# Ecology and pathogenicity for honey bee brood of recently described Paenibacillus melissococcoides and comparison with Paenibacillus dendritiformis, Paenibacillus thiaminolyticus

**DOI:** 10.1101/2025.02.06.636850

**Authors:** Florine Ory, Benjamin Dainat, Oliver Würgler, Fabian Wenger, Alexandra Roetschi, Lauriane Braillard, Jean-Daniel Charrière, Vincent Dietemann

## Abstract

Honey bee colonies contain thousands of individuals living in close proximity in a thermally homeostatic nest, creating ideal conditions for the thriving of numerous pathogens. Among the bacterial pathogens, *Paenibacillus larvae* infects larvae via the nutritive jelly that adult workers feed them, causing the highly contagious American foulbrood disease. Further *Paenibacillus* species were anecdotally found in association with honey bees, including when affected by another disease, European foulbrood (EFB). However, their pathogenicity remains largely unknown. Our results indicate that *Paenibacillus dendritiformis*, *Paenibacillus thiaminolyticus* and newly described *Paenibacillus melissococcoides* are pathogenic towards honey bee brood and that their virulence correlates with their sporulation ability, which confers them resistance to the bactericidal properties of the nutritive jelly. Our survey occasionally but increasingly detected *P. melissococcoides* in confirmed and idiopathic cases of EFB but never in healthy colonies, suggesting that this bacterium is an emerging pathogen of honey bee brood. Overall, our results suggest that virulence traits allowing a pathogenic or opportunistically pathogenic habit towards honey bee brood are frequent in *Paenibacillus spp*., but that their degree of adaptation to this host varies. Our study clarifies the ecology of this ubiquitous genus, especially when infecting honey bees.

## Introduction

The proximity of a large number of individuals makes the colonies of social insects a favorable ground for infestations by parasites and infections by pathogens (Schmid-Hempel, 1998; Cremer et al., 2018). Parasites and pathogens of the honey bee have been particularly well studied because of the important economic role these insects play, by providing crop pollination and hive products (Gallai et al., 2009; Khalifa et al., 2021). Among these pathogens, bacteria affecting honey bee brood are locally highly prevalent in several countries. *Melissococcus plutonius* and *Paenibacillus larvae*, the etiological agent of European foulbrood (EFB) and American foulbrood, respectively, are such bacteria (Hansen and Brodsgaard, 1999; Grossar et al., 2023). Outbreaks of these diseases incurs high costs to the beekeeping sector and to veterinary authorities (Genersch, 2010; Grangier et al., 2016).

Foulbroods can affect colony development and in severe cases lead to colony collapse. *M. plutonius* and *P. larvae* infect first instar larvae through midgut colonization after ingestion of contaminated nutritive jelly provided by nurse honey bees (Forsgren, 2010; Genersch, 2010). Aside *M. plutonius*, several other bacteria, including *Paenibacillus* species, are detected in EFB diseased larvae and are described as EFB secondary invaders (*Achromobacter euridice, Bacillus pumilus, Brevibacillus laterosporus, Enterococcus faecalis, Paenibacillus alvei, Paenibacillus dendritiformis*; (Forsgren, 2010; Erler et al., 2014; Gaggia et al., 2015). Each of these organisms has been suspected to be pathogenic for honey bee brood. However, their role in the course of the EFB disease is still debated (Erban et al., 2017) and apart for heat activated spores of *P. alvei* (Grossar et al., 2020), their pathogenicity to honey bee brood has not been established yet. This is also the case for *Paenibacillus thiaminolyticus* found in honey bee larvae (Nakamura, 1990; Shida et al., 1997) and *Paenibacillus melissococcoides*, a novel, fully sequenced, EFB-associated bacterial species, isolated from the nutritive jelly provided to first instar worker larvae in a EFB-diseased colony (Ory et al., 2023; Dainat et al., 2023). Because of their ecology, *Paenibacillus* bacteria seem likely to encounter honey bees and many species may possess virulence traits allowing them to behave as pathogens or opportunistic pathogens, as hypothesized in other organisms (Celandroni et al., 2016; Grady et al., 2016; Keller et al., 2018). Identifying factors that influence the virulence of pathogens is central to our ability to design prevention and control measures (Diard and Hardt, 2017).

Here, we evaluated the pathogenicity of *P. dendritiformis, P. thiaminolyticus and P. melissococcoides* for honey bee brood by oral inoculation via the nutritive jelly of first instar larvae reared *in vitro*. In addition, we investigated whether the recently described species *P. melissococcoides* could be classified as a honey bee brood pathogen by testing Koch’s postulates (Falkow, 2004; Cohen, 2017). To better understand the factors that influence the susceptibility of brood to *P. melissococcoides* infection, we conducted an age- and dose-dependence virulence assay in which we orally exposed larvae of different ages to the bacteria as well as exposed larvae to different bacterial doses. Finally, we assessed the geographic distribution and spread of *P. melissococcoides* in Switzerland by screening for its presence in symptomatic and asymptomatic EFB colonies and idiopathic EFB cases (colonies with brood disease symptoms, but without *M. plutonius* infections) sampled throughout the country between 2005 and 2021. To determine whether the bacterium was detected in other regions of the world, we screened the DDBJ/ENA/GenBank genetic sequence databases. It is indeed possible that this bacterium was previously detected elsewhere, but misidentified due to its genetic proximity to *P. dendritiformis* and *P. thiaminolyticus* (Ory et al., 2023; Dainat et al., 2023). By showing pathogenicity of several *Paenibacillus* species, including the newly described species *P. melissococcoides*, our results provide a deeper understanding of the ecology and evolutionary trajectory of the genus *Paenibacillus* when associated with honey bees.

## Results

### Ecology of newly described *Paenibacillus melissococcoides*

*P. melissococcoides* originally grew in culture together with *M. plutonius* after plating nutritive jelly collected from an EFB diseased colony (Ory et al., 2023). In the present study, we screened for the occurrence of *P. melissococcoides* and *M. plutonius* in the jelly of 40 brood cells containing 1^st^ instar larvae in the colony the bacterium was discovered in. The jelly of 30 of these 40 cells contained *P. melissococcoides* with a mean (SD) of 3.9 (4.8) CFUs per microliter of jelly (range 1 – 20), compared to four cells for *M. plutonius* with a mean (SD) of 25.3 (28.6) CFUs per microliter. The jelly of three cells contained both bacteria species. Neither of the bacteria was found in eight cells. The larvae developing in four of these 30 cells contained 1, 4, 6 and 6 CFUs of *P. melissococcoides* per microliter. In one of latter cells, the bacterium was not detected in the jelly from which the infected larva was feeding. *M. plutonius* grew from two other larvae (1 and 5 CFUs). Only in one of these cases was *M. plutonius* also found in the nutritive jelly. No bacteria grew from the remaining 36 larvae despite the presence of bacteria in the jelly some of them fed from. Because of the low number of bacteria and the complex matrices they occurred in, it was not possible to determine microscopically if the bacteria were in the vegetative or spore form.

One year after its first isolation, during a new EFB outbreak at the same source apiary, the bacterium was again detected in larvae and also in adult workers in all EFB symptomatic colonies. By contrast, *P. melissococcoides* was not detected in workers or larvae of the single asymptomatic colony on this apiary (Table S1).

### Pathogenicity towards honey bee brood of *Paenibacillus thiaminolyticus, Paenibacillus dendritiformis* and *Paenibacillus melissococcoides*

Three *P. melissococcoides* colonies, one of each of three cells/jelly from the colony it was discovered in, were picked and grown in pure cultures for our pathogenicity tests. To determine whether they were pathogenic to honey bee brood, these three isolates (labelled 1.2, 2.1, and 3.2), as well as the two genetically closely related *Paenibacillus* species, *P. dendritiformis* LMG 21716 and *P. thiaminolyticus* DSM 7262 were orally inoculated to worker larvae. The larvae were reared *in vitro* and inoculation was performed at the 1^st^ instar stage via their diet, mirroring the natural infection pathways of brood bacterial pathogens such as *P. larvae* and *M. plutonius*. *M. plutonius* CH 49.3, a highly virulent strain, was used as a positive control. Inoculation of honey bee brood with 2.10^5^ CFUs of *M. plutonius* CH 49.3, *P. melissococcoides*, *P. dendritiformis* or *P. thiaminolyticus* significantly reduced their survival probability compared to control brood (Fig. 1, Table S2). Most of the larvae inoculated with *P. melissococcoides* (71%) and *P. dendritiformis* (98%) that died, died by the fourth day of rearing. By contrast, an increased mortality of larvae inoculated with *P. thiaminolyticus* was observed at the time of pupation on the tenth day of rearing. Larvae inoculated with *M. plutonius* mostly died between the sixth and tenth day of rearing. The surviving larvae completed their development to the imaginal stage.

**Figure 1.**
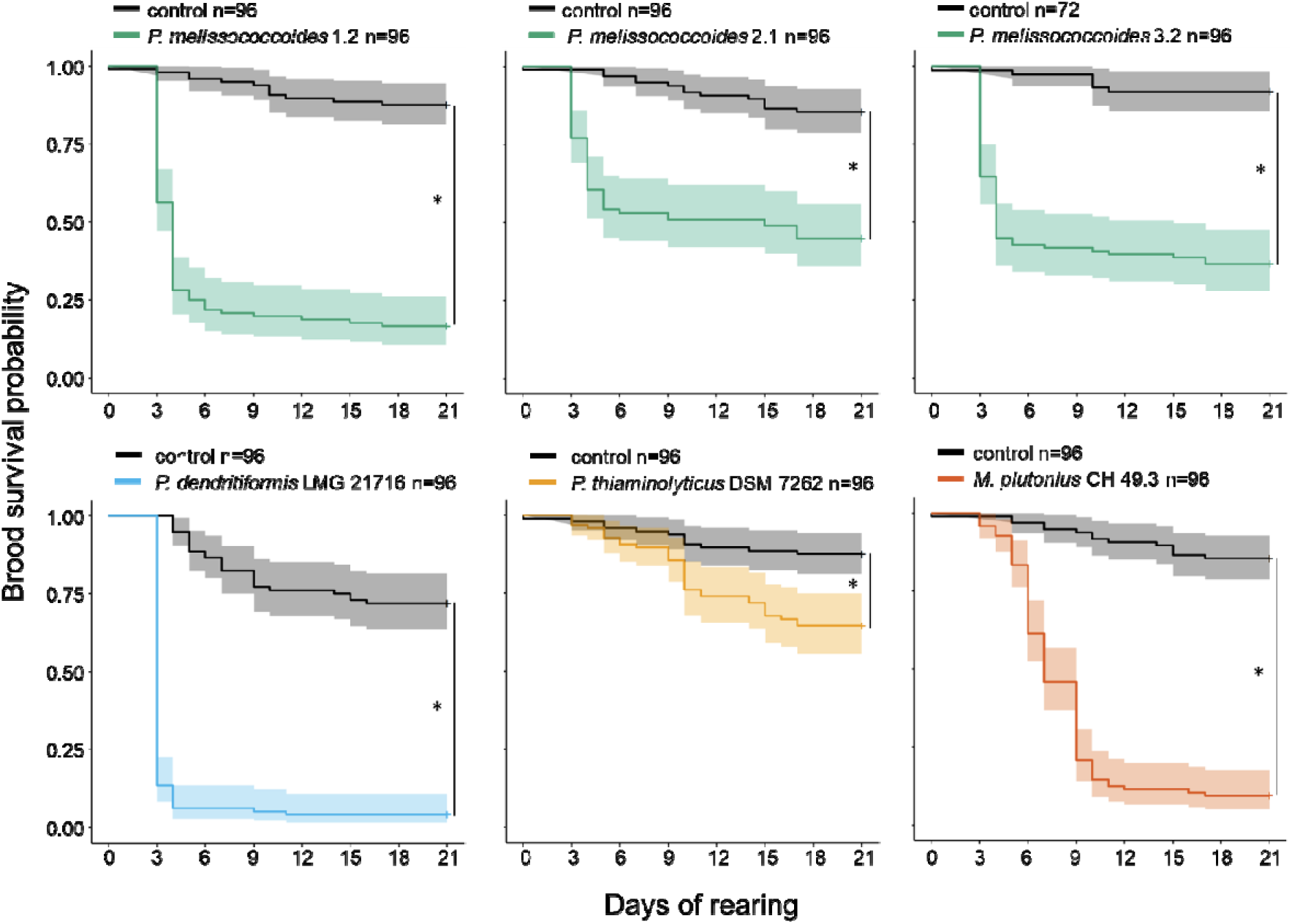
Survival profile of *in vitro* reared larvae inoculated at first instar stage with three isolates of *Paenibacillus melissococcoides,* and one isolate each of *Paenibacillus dendritiformis, Paenibacillus thiaminolyticus* and *Melissococcus plutonius*, compared with non-inoculated control larvae. Sample size is indicated in the legend. The asterisks represent significant differences between the groups’ survival curves (log-rank tests, p < 0.05, Table S2).

To assess the maximal dose ingested by the larvae, we quantified the number of *M. plutonius, P. dendritiformis, P. thiaminolyticus* and of the three isolates *of P. melissococcoides* bacteria that survived exposure to the larval diet. For this, each diet used for larval feeding was plated on agar 1.5 hours post-inoculation. No growth was observed for *P. thiaminolyticus* (Table S3), despite its presence in the bacterial suspension used to prepare the inoculum (Table S4). By contrast, CFUs were recovered from the diets containing the other bacteria with 5- to 149-fold decreases compared to the original number of bacteria mixed in the diet (Table S3). The decrease in CFUs was lower for *M. plutonius* (5.0-fold) compared to *P. dendritiformis* (9.1-fold) and *P. melissococcoides* (average decrease 59.3-fold, Table S3).

To determine which form is infections we determined the ratio of spores to vegetative cells in the bacterial stock suspensions used to prepare the inoculum. We then calculated the number of spores in three suspensions produced under the same conditions as our inocula and compared it with the number of bacteria recovered 1.5 and four hours after mixing the suspension with the diet. Indeed, the sporulation conditions of these *Paenibacillus* species are currently not known, which prevented the production of spores-only suspensions. The suspensions were composed of a mixture of vegetative cells and spores, which ratio was constant between replicates but varied among bacteria species (Table S5). The number of viable cells recovered from the diet decreased over time compared to the suspension before mixing with the diet, with CFU numbers approaching the number of spores in the suspensions (Fig. S1).

The number of CFUs recovered from larvae (N=7) experimentally infected with *P. melissococcoides* isolates 1.2 and 2.1 increased along brood development and ranged from 420 to 660 at the stage L3, 60 to 4572 at L4 and >2.2×10^4^ at L5 (Table S6).

### Verifying Koch’s postulates to confirm the pathogenicity of Paenibacillus melissococcoides

To investigate the association between the occurrence of *P. melissococcoides* and brood disease, honey bee samples (N=414, including pools of several colonies) were collected from 939 colonies in 287 apiaries. EFB symptoms were observed in 786 colonies of 262 apiaries. Four colonies in four apiaries showed brood disease symptoms but were negative for *M. plutonius* (in one of these idiopathic cases the presence of *P. larvae* was excluded by microscopy) and 153 colonies in 41 apiaries were asymptomatic. Samples from individual colonies or from pools of colonies were screened for *P. melissococcoides* and *M. plutonius* by qPCR. Neither *M. plutonius* nor *P. melissococcoides* were detected in asymptomatic colonies in the period 2005-2021 (Table S7). Excepted the four idiopathic cases, *M. plutonius* was detected in all EFB symptomatic colonies in the period 2005-2020 and in all but two colonies in 2021 (Table S7). This result for 2021 is in line with the 2005-2020 samples originating from *M. plutonius* colonies previously verified as positive by qPCR, and with the 2021 samples originating from colonies previously identified as symptomatic only based on visual inspection alone and thus without a PCR confirmation of *M. plutonius* presence. *P. melissococcoides* was absent from samples (N=201) of EFB symptomatic colonies (N=617) collected between 2005 and 2010 (Fig. 2, Table S7), but was detected in one out of 84 samples from symptomatic colonies collected from 78 apiaries in 2013, one of 4 colonies and apiaries in 2014, two out of four colonies and apiaries in 2015, two out of three colonies and apiaries in 2020 and three out of 70 samples collected from 69 apiaries in 2021 (Fig. 2, Table S7). *P. melissococcoides* was detected in two of the four colonies showing symptoms of brood diseases, but in which *M. plutonius* (and *P. larvae* in two of these) were not detected, i.e. idiopathic cases (Table S7).

**Figure 2.**
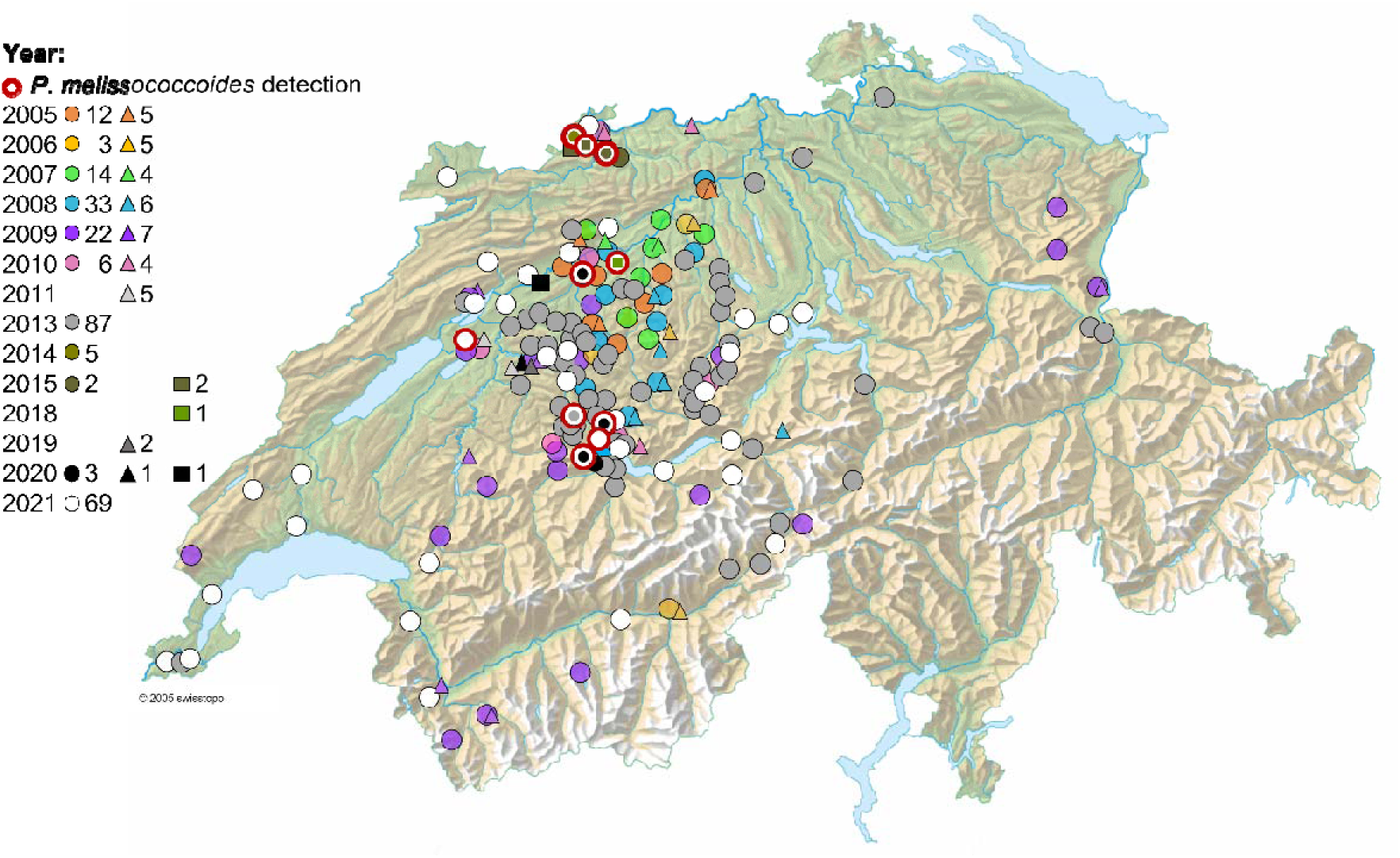
Location of apiaries screened for *Paenibacillus melissococcoides* in the period 2005-2021 in Switzerland. Number of apiaries analyzed per year and symptom status is indicated next to the corresponding symbol in the legend: circles correspond to screened apiaries with EFB-symptomatic colonies, triangles to screened apiaries with asymptomatic colonies and squares to idiopathic cases. Colors represent the different years. In many occasions, apiaries screened were so close to each other that their symbols overlap completely.

To investigate pathogenicity at the individual level of immature honey bees, *P. melissococcoides* originating from the pure cultures of the three isolates 1.2, 2.1, and 3.2 (Fig. 3A, step 2) were inoculated to honey bee larvae reared *in vitro*. Exposure to *P. melissococcoides* significantly enhanced larval mortality compared to non-inoculated controls (Fig. 1, Fig. 3A, step 3). Despite the presence of this strain in the data base, *P. melissococcoides* 1.2 reisolated from the inoculated dead brood reared *in vitro* was identified as being identical to *P. melissococcoides* 3.2 by protein profile analysis with MALDI-TOF mass-spectrometry (Fig. 3A). This reisolate again caused mortality after inoculation of healthy larvae (Fig. 3A, step 4, Fig. 3B).

**Figure 3.**
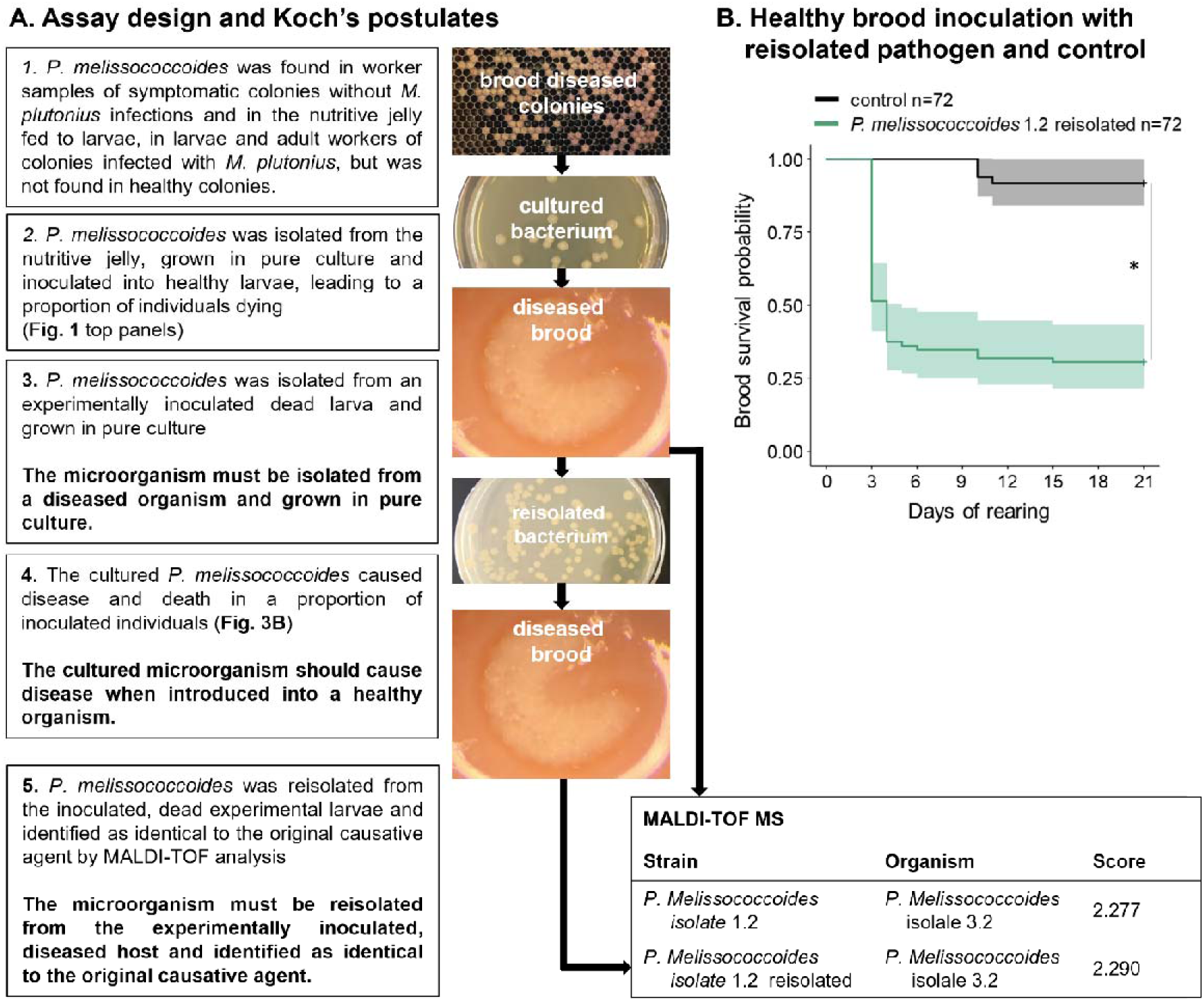
*Paenibacillus melissococcoides* pathogenicity to honey bee brood. **A.** Assay design and corresponding steps of the Koch’s postulates (in bold) testing the pathogenicity of *P. melissococcoides* at the individual level. Also shown are results of MALDI-TOF mass-spectrometry identification (column ‘organism’) of *P. melissococcoides* after sequential isolation and re-isolation from larvae that died following inoculation with isolate 1.2 (column ‘strain’). Score value interpretation: 3.000-2.300: highly probable species identification; 2.299-2.000: secure genus identification, probable species identification; 1.999-1.700: probable genus identification; 1.699-0.000: unreliable identification. **B.** Survival profile of *in vitro* reared larvae inoculated at first instar stage with *P. melissococcoides* 1.2, which was reisolated from a previously inoculated dead fifth instar larva. Survival was compared with non-inoculated control larvae. Sample size is indicated in the legend. The asterisk represents a significant difference between the groups’ survival curves (log-rank test).

### Effect of inoculum dose and host age on larval susceptibility to Paenibacillus melissococcoides

Honey bee brood showed a dose- and age-dependent susceptibility to *P. melissococcoides* exposure. The brood survival probability significantly decreased with increasing dose of bacterial cells in the inoculum fed to the larvae, and their susceptibility to inoculation significantly decreased with increasing age (Fig. 4, Tables S8-S9).

**Figure 4.**
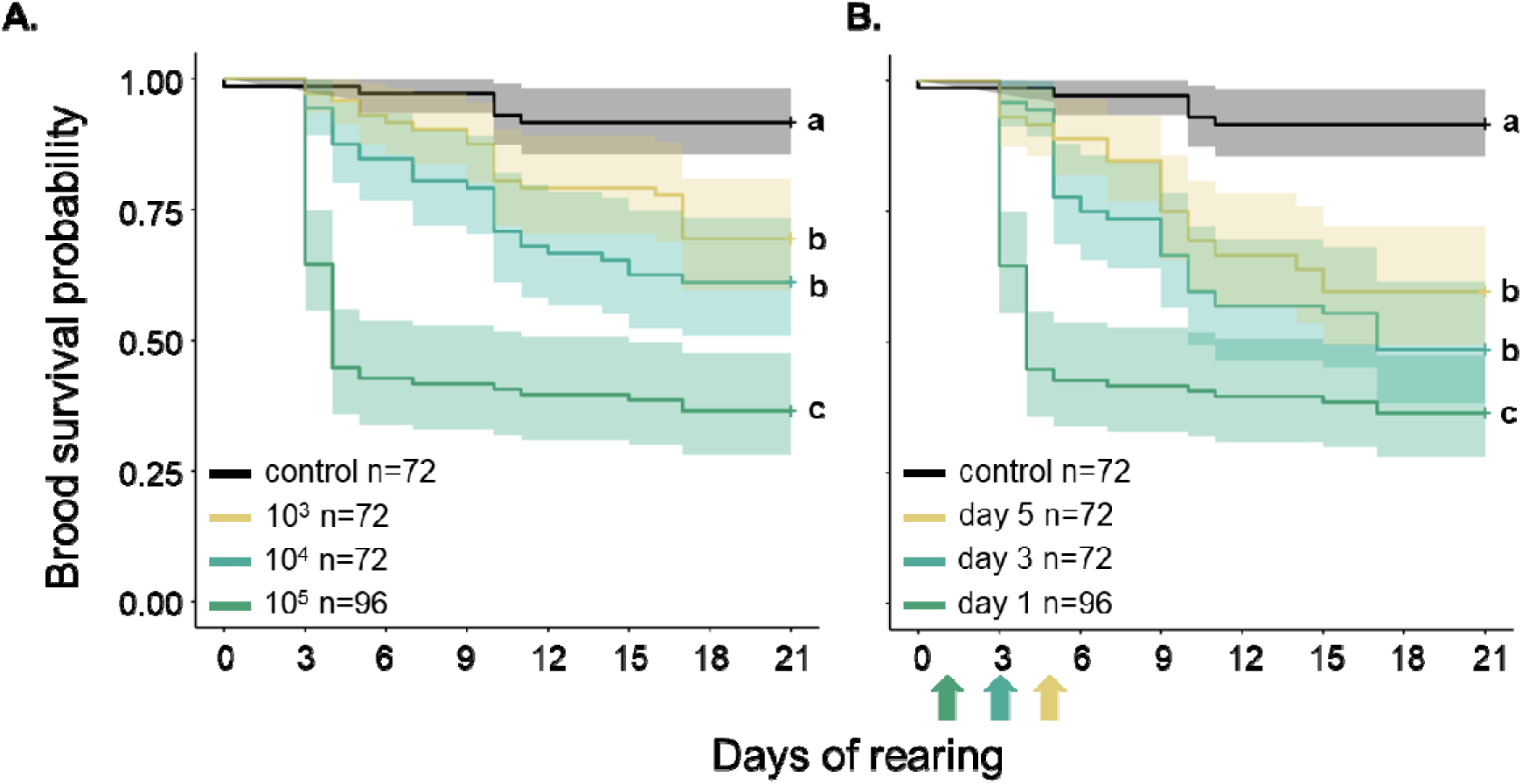
Survival profiles of *in vitro* reared larvae inoculated with *Paenibacillus melissococcoides* 3.2 and of non-inoculated control larvae. **A. Dose dependence:** one-day old larvae inoculated the first day of rearing with different doses of *P. melissococcoides* (in CFUs) and controls. **B. Age dependence:** one-, three- and five-day old larvae inoculated with 10^5^ *P. melissococcoides* bacteria (arrows), respectively, and controls. Sample size is indicated in the legend. Different letters to the right of the curves indicate significant differences in the survival of the tested dose and age groups (pairwise log-rank tests, Bonferroni-Holm corrected, p<0.05).

### Geographic distribution of *Paenibacillus melissococcoides*

To investigate the distribution area and spread of *P. melissococcoides* in Switzerland, honey bee samples (N=414, including pools of several colonies) were collected in the period 2005-2021 from 786 colonies showing EFB symptoms in 262 apiaries, from four colonies showing brood disease symptoms in absence of *M. plutonius* (and *P. larvae* in two of them) in four apiaries and from 153 asymptomatic colonies in 41 apiaries. *P. melissococcoides* was detected in a 2013 sample collected approximately 10 km from its discovery location seven years later, in 2020. *P. melissococcoides* was again found in 2014, 2015, 2018 and 2020 samples of five apiaries distant of up to 90 km of its discovery location. In 2021, the bacterium was found again at the site of discovery (Ory et al., 2023) as well as in two more apiaries, 8 and 75 km away.

To determine whether the bacterium was detected in other regions of the world, but possibly misidentified, the complete 16S rDNA sequence of *P. melissococcoides* isolate 2.1 was subjected to a BLAST search against the 16S rDNA sequences of *Paenibacillus spp.* (taxid: 44249) present in the DDBJ/ENA/GenBank databases. The *Paenibacillus* spp. isolates most closely related (i.e. with sequence identity >99%) to *P. melissococcoides* were *P. dendritiformis* strains isolated from alive and dead honey bee larvae in Italy (Table 1). They were followed by *P. dendritiformis* strains isolated from Japanese honey (Table 1). Genetically highly similar bacteria (previously identified as *Paenibacillus popilliae* and undescribed strains) were also found in soil, water, plant and organic waste material (Table 1).

**Table 1.** Summary table of blastN search against the 16S rDNA sequences of *Paenibacillus* spp. available in the DDBJ/ENA/GenBank databases.

| Country | Isolation source | Authors | Species/strain/isolate | Identity (%) | Accession number |
| --- | --- | --- | --- | --- | --- |
| Switzerland | Honey bee nutritive jelly | Ory et al., 2023 | *P. melissococcoides* 3.2 | 100 | OW961638.1 |
| Switzerland | Honey bee nutritive jelly | Ory et al., 2023 | *P. melissococcoides* 2.1 | 100 | OW961637.1 |
| Switzerland | Honey bee nutritive jelly | Ory et al., 2023 | *P. melissococcoides* 1.2 | 100 | OW961636.1 |
| Italy | Honey bee | Gaggia et al., | *P. dendritiformis* strain | 99.93 | KR073927.1 |
|  | larvae | 2015 | NDHL-P1 |  |  |
| Italy | Dead honey bee larvae | Alberoni et al., 2015 (unpublished) | *P. dendritiformis* strain PA(C) | 99.93 | MG650035.1 |
| Italy | Dead honey bee larvae | Alberoni et al., 2015 (unpublished) | *P. dendritiformis* strain PA(B) | 99.93 | MG650034.1 |
| Japan | Japanese honey | Okamoto et al., 2021 | *P. dendritiformis* J47TS5 | 99.87 | LC588610.1 |
| Japan | Japanese honey | Okamoto et al., 2021 | *P. dendritiformis* J46TS7 | 99.87 | LC588605.1 |
| Japan | Japanese honey | Okamoto et al., 2021 | *P. dendritiformis* J44TS4 | 99.87 | LC588592.1 |
| Italy | Honey bee larvae | Gaggia et al., 2015 | *P. dendritiformis* strain SDHL-P7 | 99.86 | KR073928.1 |
| Japan | Japanese honey | Okamoto et al., 2021 | *P. dendritiformis* J22TS4 | 99.8 | LC588505.1 |
| Japan | Japanese honey | Okamoto et al., 2021 | *P. dendritiformis* J13TS5 | 99.8 | LC588473.1 |
| Japan | Japanese honey | Okamoto et al., 2021 | *P. dendritiformis* J11TS3 | 99.8 | LC588468.1 |
| Japan | Japanese honey | Okamoto et al., 2021 | *P. dendritiformis* J10TS2 | 99.73 | LC588466.1 |
| India | Water | Nandanwar, 2018 (unpublished) | *P. dendritiformis* strain PV3-16 | 99.68 | MH472941.1 |
| Japan | Japanese honey | Okamoto et al., 2021 | *P. dendritiformis* J4TS5 | 99.67 | LC588445.1 |
| India | Bird feces | Joshi et al., 2013 (unpublished) | *Paenibacillus* sp. BAB-3420 | 99.66 | KF917150.1 |
| India | Bird feces | Joshi et al., 2013 (unpublished) | *Paenibacillus* sp. BAB-3408 | 99.66 | KF917139.1 |
| India | na | Khan, 2015 (unpublished) | *P. dendritiformis* strain ANSK05 | 99.66 | KT152690.1 |
| India | Bird feces | Joshi et al., 2013 (unpublished) | *Paenibacillus* sp. BAB-3421 | 99.66 | KF917151.1 |
| China | Congo Red | Gan, 2017 | *P. dendritiformis* strain | 99.65 | KY655213.1 |
|  | dye | (unpublished) | GGJ7 |  |  |
| Soudan | Soil | Gorashi et al., 2012 (unpublished) | *P. popilliae* strain Sh-14 | 99.64 | KC201676.1 |
| Japan | Japanese honey | Okamoto et al., 2021 | *P. dendritiformis* J8TS3 | 99.6 | LC588463.1 |
| Japan | Japanese honey | Okamoto et al., 2021 | *P. dendritiformis* J7TS5 | 99.6 | LC588460.1 |
| Japan | Japanese honey | Okamoto et al., 2021 | *P. dendritiformis* J6TS7 | 99.6 | LC588455.1 |
| Japan | Japanese honey | Okamoto et al., 2021 | *P. dendritiformis* J3TS5 | 99.6 | LC588442.1 |
| Japan | Japanese honey | Okamoto et al., 2021 | *P. dendritiformis* J1TS6 | 99.6 | LC588431.1 |
| India | Kodo millet | Parthiban, 2016 (unpublished) | *P. dendritiformis* strain PP | 99.6 | KX082752.1 |
| Pakistan | na | Bhutto and Bhutto, 2022 (unpublished) | *P. dendritiformis* strain IBGE-MAB1 | 99.6 | OP648144.1 |
| France | T168 culture | Roux and Raoult, 2003 (unpublished) | *P. dendritiformis* strain T168 | 99.59 | NR_042861.1 |

## Discussion

Inoculations with *P. thiaminolyticus, P. dendritiformis and P. melissococcoides* led to significantly decreased survival of honey bee brood compared to non-inoculated controls. Pathogenicity towards the brood was confirmed for the novel species *P. melissococcoides* by fulfilling three of four of Koch’s postulates at the individual level and by showing the multiplication of the bacteria in infected diseased hosts. Our results indicate that spores are the infectious form. At the colony level in the field, *P. melissococcoides* was not detected in 153 healthy colonies but in two diseased colonies negative for *M. plutonius*, of which one was also negative for *P. larvae*, bringing some support to the remaining Koch postulate. *P. melissococcoides-*induced host mortality increased with inoculum dose and decreased with host age. Between 2013 and 2021, this bacterium was found in 10 out of 177 brood diseased apiaries in Switzerland, but was not detected in older samples from EFB-diseased apiaries (2005-2010, N = 185) or from asymptomatic colonies (2005-2021, N = 153). Even though the distribution area of *P. melissococcoides* appears limited to a small area of Switzerland, it possibly extends beyond the country’s borders because isolates with highly similar sequences deposited in repositories were found in Italy and Japan (Table 1).

Inoculation of first instar honey bee larvae with the three bacteria species studied decreased their survival. The mortality due to *P. dendritiformis* inoculation, to a level comparable to that of highly virulent *M. plutonius* CH 49.3, was higher compared to the two other species. The difference in virulence observed is in line with the survivability of these bacteria in the nutritive jelly fed to larvae. The number of CFUs recovered after 1.5 h exposure to larval diet (Table S3) correlated with brood mortality rates they generated (Fig. 1). Viable *M. plutonius* bacteria were recovered in the largest number, followed by *P. dendritiformis* and *P. melissococcoides,* whereas *P. thiaminolyticus* did not grow at all, despite the presence of viable bacteria in the suspension used for inoculum preparation (Table S4). The capacity of bacteria to survive an exposure to the jelly is thus likely an important virulence factor (Takamatsu et al., 2017; de La Harpe et al., 2022) and was here correlated to the number of spores present in the stock suspension (Table S5, Fig. S1). Our results indicate that vegetative cells perished rapidly in the diet and that spores survived (Fig. S1) to later germinate in the larvae. As for *P. larvae*, spores of the tested bacteria thus appear to be the infectious form. A biologically more relevant assessment and comparison of the virulence of the tested bacteria should be obtained from *in vitro* tests using nutritive jelly as fed by nurse bees instead of experimental diet (de La Harpe et al., 2022) and equalized number of spores among *Panibacillus* spp. inocula devoid of vegetative cells.

The pathogenicity of *P. melissococcoides* for honey bee brood was confirmed by the fulfillment of three of the four Koch’s postulates (Fig. 3) applied at the individual level and with some support to the remaining postulate at colony level. The decrease in brood survival following exposure to *P. melissococcoides* was not due to a toxic effect of dead bacteria in the diet because the number of bacteria increased during larval development to the point of surpassing the number of bacteria in the inoculum (Table S6). We can also safely exclude a saprophytic mode of action because inoculation with pure *P. melissococcoides* suspensions led to larval death (Fig. 1, 3B). As for *M. plutonius* and *P. larvae* (Forsgren et al., 2018), young larvae were more susceptible to *P. melissococcoides* infection than older ones (Fig. 4B). Also, similarly to *P. larvae* (Hernández López et al., 2014), inoculation with an increasing number of *P. melissococcoides* bacteria led to an increase in mortality (Fig. 4A).

The exposure pathway and number of bacteria used in our *in vitro* assays were biologically relevant for *P. melissococcoides*. The bacterium was found in the nutritive jelly and larvae in the colony from which it was isolated for the first time. Based on Thrasyvoulou and Benton (1982), we estimated that a larva of 24 to 48h of age (as used in our assays) ingests 1 mg of jelly. In the colony in which *P. melissococcoides* was discovered, 1 mg of jelly contained a mean of 3.9×10^3^ (SD 4.8×10^3^, range 1×10^3^ – 2×10^4^) CFUs, likely originating from spores because of the low survival of vegetative cells in this medium (Fig. S1). This range corresponded well to the number of spores present in our inocula (2.4×10^2^ – 2.4×10^4^, calculated from the average 12% spores out of the 2×10^3^ to 2×10^5^ CFUs, i.e. a mixture of spores and vegetative cells, in our inocula, Table S5).

*P. melissococcoides* was also found on adult workers, which probably function as transmission vectors of the bacterium among larvae. We were not able to determine whether workers carried vegetative cells or spores. The biology of *P. melissococcoides* requires further investigation to determine sporulation dynamics in the hive environment and to explain its presence at higher prevalence compared to *M. plutonius* (i.e. proportion of the 40 cells screened in the EFB diseased colony, 30 vs. 4 cells, respectively).

*P. melissococcoides* was pathogenic for individual immatures and not found in healthy colonies (N = 153). We also found the bacterium twice in honey bee colonies showing symptoms of a brood disease while negative for *M. plutonius* (and *P. larvae* in at least one of these cases, Table S7). Because we could not safely exclude the presence of further pathogens in these colonies, we could not fulfill with high confidence the first half of Koch’s postulates stating that the microorganism must be found in abundance in all organisms (colonies) suffering from the disease. It is thus not established whether *P. melissococcoides* can trigger a disease at the colony level on its own and a large screening of colonies showing brood diseases for *P. melissococcoides* or experimental infections of colonies are required. Even if *P. melissococcoides* did not need the presence of *M. plutonius* to cause mortality of *in vitro* reared honey bee brood, this lack of clear association with clinical cases positive for *P. melissococcoides* only, currently places this bacterium in the group of secondary invaders associated with EFB.

Between 2013 and 2021, *P. melissococcoides* was detected in apiaries within a 30×90 km region. In 2021, the bacterium was found again at the site of discovery (Ory et al., 2023). Transmission routes of *P. melissococcoides* are unknown to date, but can include natural and anthropic processes via drifting of foragers, robbing of infected colonies by others, exchange of infected beekeeping material (as is the case for *M. plutonius* and *P. larvae*, Forsgren, 2010; Genersch, 2010), other insects (Deutsch et al. 2023) or yet unidentified vectors.

*P. melissococcoides* was thus found repeatedly and increasingly in samples dating back to 2013 but not in samples collected before this year and as far back as 2005, and this despite a similar sampling effort across the two periods. This pathogen could thus be emerging or alternatively have been recently imported from elsewhere. Indeed, we found possible occurrence of this bacterium in EFB-diseased colonies beyond Switzerland. The cases described in the north and south of Italy were genetically highly similar to *P. melissococcoides* (Table 1), although the Italian isolates were originally ascribed to *P. dendritiformis* (Gaggia et al., 2015). A bacterium with very high genetic similarity to *P. melissococcoides* was also found in Japanese honey samples (Okamoto et al., 2021). In addition, we found possible occurrence of *P. melissococcoides* misidentified as *P. dendritiformis* in various environments beyond the honey bee colony (Table 1), which suggests that *P. melissococcoides* could be widespread, as is typical for the genus *Paenibacillus* (Grady et al., 2016). There is a need for screenings at larger scale to better understand the distribution area and population dynamics of this bacterium, as is required for other honey bee brood pathogens (Grossar et al., 2023).

*Paenibacillus* species are ubiquitous in the environment and many are tightly associated with plants (Grady et al., 2016). Their presence on plants, which the honey bees visit to collect their food, can result in frequent interactions, a prerequisite to develop an association with this host. Their abilities to survive in the nutritive jelly, which honey bee larvae must ingest for the bacteria to reach their replication milieu (e.g. de La Harpe et al., 2022) appears to depend on their ability to sporulate. Because of their pathogenic effect on individual immatures, there is potential for *P. melissococcoides*, *P. dendritiformis* and *P. thiaminolyticus* to cause disease at the colony level on their own, although other virulence factors might be required to trigger a pathology and no such case has been reported to date. The ability of *Paenibacillus* species to exploit and damage honey bee brood can be facilitated by the high number of enzymes they are known to produce. Enzymes may allow them to outcompete other species of the gut microbiome or to attach to and damage the peritrophic matrix to gain entry into the midgut epithelium and hemocoele (Genersch, 2010; Grady et al., 2016). Because the pathogenic *Paenibacillus* species are phylogenetically distant from each other (Ory et al., 2023), it appears that the genus commonly possesses virulence traits to allow pathogenic or opportunistic pathogenic habits in a large range of hosts (Celandroni et al., 2016; Grady et al., 2016; Keller et al., 2018). This idea is supported if *P. melissococcoides* and not *P. dendritiformis* was found in Italian and Japanese samples (Gaggia et al., 2015; Okamoto et al., 2021), suggesting that *P. dendritiformis* was to date never associated to honey bees (see also Erban et al. 2017) but nevertheless showed a high virulence towards their brood. The genus *Paenibacillus* thus represents a good model system to investigate the determinants of pathogenicity and virulence in bacteria (Diard and Hardt, 2017).

## Conclusion

Our results indicate that *P. melissococcoides* is pathogenic to honey bee brood and a secondary invader associated with EFB. There is as yet no strong evidence that *P. melissococcoides* triggers a disease at the colony level on its own, but we found the bacterium in idiopathic cases, in which symptoms of brood diseases are detected without the presence of *M. plutonius* or *P. larvae* in the affected colony (Shimanuki and Knox, 1994; vanEngelsdorp et al., 2013, this study), suggesting that pathogenicity of *P. melissococcoides* at the colony level is possible. *P. melissococcoides* has to date not been found outside honey bee colonies (but see Table 1) and whether it is an obligate or facultative pathogen of honey bees is not known. Overall, the genus *Paenibacillus* seems to commonly be able to express a pathogenic habit in the honey bee brood, but the degree to which each species is likely and adapted to do so remains to be determined.

## Material and method

### Bacterial strains/isolates and cultivation

Three *P. melissococcoides* isolates, 1.2, 2.1 and 3.2 (Ory et al., 2023), and two closely related *Paenibacillus* species, *P. dendritiformis* LMG 21716 (type strain, provided by BCCM LMG) and *P. thiaminolyticus* DSM 7262 (type strain, provided by DSMZ GmbH) were cultivated on basal medium Petri dishes for four days at 36°C under oxic conditions as recommended in Ory et al. (2023). The basal medium contained 10 g/l yeast extract, 10 g/l glucose, 10 g/l starch, 20 g/l agar, 0.25 g/l L-cysteine and 100 ml 1M KH_2_PO_4_ in distilled water, adjusted to pH 6.7 using 2.5M KOH and autoclaved at 115°C for 15 minutes (Budge et al., 2024). *M. plutonius* strain CH49.3 was chosen as positive control because of its high virulence (Grossar et al., 2020). *M. plutonius* CH49.3 was also cultivated on basal medium at 36°C but for five days and under anoxic condition (hermetic box with anoxic generator sachet, GENbox anaer, bioMérieux and anoxic indicator) as described in Budge et al. (2024).

After incubation, the bacterial colonies that grew on the Petri dishes were suspended into liquid basal medium. An aliquot of this suspension was used to determine its concentration in colony forming units (CFUs) and the rest was stored at 4°C until use to prepare the inocula. To determine CFU concentration, ten-fold serial dilutions of the aliquot were prepared and cultivated on basal medium Petri dishes for four days as explained above. CFUs on three Petri dishes were then counted and the counts averaged (Budge et al., 2024). After the four days, the suspensions were adjusted by dilution with sterile saline buffer to the concentration desired for the inocula (see next section). Bacterial suspension and larval diet were then mixed to produce the inocula. For each inoculation experiment, a new bacterial suspension was prepared from the stock stored in glycerol at −80°C. Stock suspensions thus never remained stored for more than four days before use. Sugar and royal jelly composing larval diet are known to have antibacterial properties (Selwyn and Durodie, 1985; Molan, 1992; Erler et al., 2014; Vezeteu et al., 2016). Thus, to estimate the number of viable cells fed to larvae for inoculation assays (Lewkowski and Erler, 2018), bacterial inocula were plated after each experimental inoculation session. For this, twenty microliters of diluted bacterial inocula (1:10^2^, 1:10^3^, 1:10^4^) as well as control diet (dilution 1:10^2^) were plated 1.5 h (range 1-2h) post-inoculation on Petri dishes. CFUs were counted on three replicates and averaged after four days incubation and are given as estimated CFUs per larva.

To investigate which bacterial form was infectious for larvae and because the sporulation conditions are not known for the three bacterial species tested, we determined the ratio of vegetative cells vs. spores in suspensions produced under the same conditions as our inocula and just before mixing in the diet (3 replicates). For this, a 1:100 or 1:1000 dilution of the suspension was added to a hemocytometer to count vegetative cells and spores under a phase contrast microscope following standard methods (Human et al. 2013). In addition, aliquots of the suspension before mixing with the diet as well of the inoculum 1.5h and 4h after mixing was plated as described above. Our expectations, based on work with *P. larvae* (Hornitzky 1998), was that vegetative cells present in the suspension would rapidly die in the diet (even before having the opportunity to sporulate, which takes several hours to several days, e.g. Baril et al. 2012) due to the antibiotic properties of the royal jelly and that only spores formed before mixing with the diet would survive to germinate in the larvae. CFUs growing on the plates should thus correspond to the number of spores measured in the stock suspension, indicating that only the spores are infectious, as for *P. larvae* (Hornitzky 1998).

### Larval inoculation assays

Honey bee larvae originating from six healthy colonies of *A. mellifera* were used for larval inoculation assays. First instar worker larvae were obtained by caging the queens for 24 hours on empty combs using excluder cages (Human et al., 2013). Freshly hatched larvae were reared following previously described protocols (Crailsheim et al., 2013; Ory et al., 2022). For larval inoculation assay, the grafted larvae were deposited into 10 µl of diet A (50% pure royal jelly mixed with 50% filtered sugar (0.2 µm) prepared with pure water containing 12% D-glucose, 12% D-fructose and 2% yeast extract). Within two hours after grafting, larvae received an additional 10 µl of diet A containing 2×10^7^ CFUs ml^-1^ (9 parts diet A for 1 part bacterial suspension) or a control diet A without bacteria (9 parts diet A for 1 part suspension buffer). Hence, each inoculated larva was fed 20 µl diet A containing 2×10^5^ CFUs. On day three, inoculated and non-inoculated larvae were fed 20 µl of diet B (50% pure royal jelly mixed with 50% filtered sugar made of pure water containing 15% D-glucose, 15% D-fructose and 3% yeast extract). On days 4, 5 and 6, larvae received 30, 40 and 50 µl of diet C (50% pure royal jelly mixed with 50% filtered sugar made of ultra-pure water containing 18% D-glucose, 18% D-fructose and 4% yeast extract), respectively. The larvae were kept in an incubator at 34.5 °C and 95% relative humidity except during feeding bouts, which were performed at room temperature. For pupal development, the brood was kept at 34.5 °C and 75 % relative humidity. The royal jelly needed for larval rearing was harvested from healthy queenless colonies from our local apiary and stored at −25°C in a sterile environment until use.

To verify further steps of Koch’s postulates (Fig. 3), *P. melissococcoides* was reisolated from an *in vitro* reared larva (see previous paragraph) which died after inoculation with *P. melissococcoides* 1.2. The reisolated bacteria were also used for molecular diagnostics to confirm species identity and for a new series of larval inoculation to evaluate the maintenance of pathogenicity after passage in a larva. For this, the bacteria was reisolated by homogenizing a dead inoculated larva in 1 ml saline buffer and plating 50 µl of this soup on basal medium Petri dishes and incubating under conditions described above for *Paenibacillus* spp. growth. Plates were visually inspected for colonies showing *P. melissococcoides* morphology (Ory et al., 2023), which were then picked for bacterial identity confirmation through MALDI-TOF MS (Brucker) before the new series of larval inoculation. To determine the dose-dependence of *P. melissococcoides* virulence towards honey bee brood, groups of first instar larvae were exposed during the first day of the *in vitro* rearing to 2×10^3^, 2×10^4^, or 2×10^5^ CFUs of *P. melissococcoides* 3.2. To determine the age at which honey bee larvae are most susceptible to *P. melissococcoides*, one-, three- and five-day old larvae were exposed to 2×10^5^ CFUs of *P. melissococcoides* 3.2, corresponding to the first, third and fifth larval instars (all were grafted when one day old). Larvae of each group received a single bacterial inoculation. Control larvae were reared in the same manner as inoculated larvae except for the lack of bacteria in their diet. Brood survival was monitored daily from day 3 until completion of the development (i.e., until the imaginal stage). The brood status (dead or alive) was recorded every 24 hours until pupation was completed (day 13), by observing larvae under binocular microscope. A larva was considered dead when neither respiratory movements nor reaction to a mechanical stimulus applied with a sterile plastic needle was detected. The dead larvae were removed from the plates. Pupae are mostly immobile and respiration is difficult to observe, thus the status of the pupated individuals was determined at the end of the experiment, when live imagos emerged. Experiments were performed with a minimum of 72 larvae per group and with at least two grafting series at seven days interval. Each series consisted in a sample of 24-48 larvae per group (Fig. 1, Table S1).

To determine whether brood mortality was due to the toxicity of dead bacteria in the inoculum or live pathogenic bacteria multiplying in their host, one to two larvae at stage L3, L4 and L5 infected with *P. melissococcoides* isolates 1.2 and 2.1 were washed in 400 µl saline buffer to remove adhering jelly and crushed by vortexing in 600 µl saline buffer. Fifty microliter of the suspension was plated on basal medium as described above.

### Detection of *P. melissococcoides* in nutritive jelly and in larvae

To test for the natural occurrence of *P. melissoccoides* in the nutritive jelly fed to young worker larvae in the colony the bacterium was discovered in, 1 µl jelly from 40 cells containing 1^st^ instar larvae was collected using an inoculation loop. The microliter of jelly was mixed in 150 µl of sterile saline buffer (0.9% NaCl) and fifty microliters of this suspension were spread on each of three Petri dishes containing basal medium. CFUs were counted and averaged after four days of culture as described above. Larvae were washed in 200µl NA to exclude any bacterium in the jelly adhering to their surface, placed in 100µl NA and homogenized using a sterile needle. The homogenate was plated and CFUs counted.

### qPCR assay for *P. melissococcoides* detection

To evaluate the presence of *P. melissococcoides* in honey bee samples in Switzerland, we established a qPCR assay based on the *sodA* gene, which codes for the manganese-dependent superoxide dismutase protein. The whole genome sequences of *P. melissococcoides* isolate 2.1 (accession number, GCA_944800085) was used as reference. A 58-bp fragment with PM-F primers and Taqman^®^ MGB probe (Table S10) were designed on the *sodA* gene by using the Primer Express^®^ software (version 3.0.1, Applied Biosystems, USA). Primers and probe were purchased from Microsynth (Balgach, Switzerland).

The Takyon^TM^ No Rox 2x MasterMix UNG (Eurogentec, Belgium) was used for qPCR measurements. The qPCR was carried out in a final reaction volume of 12 µl containing 0.3 µM of each primer, 0.1 µM probe, 2 x reaction buffer and 2 µl of DNA. Amplifications were run in a Corbett Rotor-Gene 3000 using the following program: 2 min at 50°C, 3 min at 95°C, and 40 cycles of 3 s at 95°C and 20 s at 60°C. Quantitative data were analyzed using the Rotor-Gene software version 1.7.87 (Qiagen) using a threshold of 0.03 for the quantification cycle (C_q_) value determination. Each sample was measured in duplicate and non-template controls tested negative in all qPCR runs. A plasmid for the standard curve was constructed by inserting a part of the *sodA* gene into a pGEM^®^-t easy Vector System I (Promega, Switzerland) according the procedure described in Moser et al. (2017). Specificity of the qPCR assay was assessed with DNA from the following species: *P. larvae* (DSM 7030 and a wild strain, 87), *P. alvei* (DSM 29 and two wild strains, 49.1, 90), *P. dendritiformis* (LMG 21716) and *P. thiaminolyticus* (DSM 7262). Samples (12 g of worker bees from the brood nest) were homogenized and extracted according to Roetschi et al. (2008) except that a bead homogenization procedure with zirconia beads was used to extract DNA from the pellet. Samples were considered as negative when no signal (Cq = N/A, no answer) was detected at the end of the amplification and positive when averaged Cq ≤ 55. This Cq threshold was chosen due to the lack of detection limit for *P. melissococcoides* and to allow for a high detection level. Only two Cq values for *M. plutonius* were marginally above the more usual 40 threshold value and none for *P. melissococcoides*.

### Geographic distribution of *P. melissococcoides*

To determine the geographic distribution of *P. melissococcoides*, we screened samples collected between 2005 and 2021 from *M. plutonius* negative asymptomatic colonies, from *M. plutonius* positive samples from EFB-diseased colonies, as well as from colonies showing brood disease symptoms but without *M. plutonius* infection. In four cases, the absence of *P. larvae* could be confirmed microscopically. In all the remaining cases, the presence of *P. larvae* could be safely excluded by the field visual diagnostics by trained apiary inspectors. We detected, by PCR or microscopy, only one confusion between the symptoms of both diseases in 77 cases (data not shown). The 2005-2020 samples (N=331, Table S7) consisted of pools of 7 to 97g of workers collected from the brood nest of one or more of 856 colonies from 218 Swiss apiaries. These samples were previously found positive or negative for the presence of *M. plutonius* bacterium by qPCR following the method described in Roetschi et al. (2008). The extracted DNA was kept at −25°C and reused for the present study. To obtain 2021 samples, we requested 1-4 suspicious larvae from brood combs of colonies showing EFB symptoms from an official diagnostics laboratory involved in the EFB diagnostics in the field. We obtained 70 samples from 69 apiaries (Table S7). These larvae were pooled per colony and stored at −25°C until DNA extraction. For extraction, larvae were thawed, supplemented with 1 ml sterile saline (0.9% NaCl) and homogenized using sterile pestles. In a pre-lysis step, 100 µl of homogenate were transferred into 200 µl of a lysozyme solution (20 mg lysozyme/ml in: 20 mM Tris/HCL, 2 mM EDTA, 1% Triton X-100, pH=8) and incubated one hour at 37°C. Further lysis and DNA extraction were performed according to manufacturer’s protocol (NucleoSpin Tissue kit, Macherey-Nagel). The qPCR was set up in a 20 µl volume, including 2 µl DNA-extract volume with Master Mix (2x) Universal (KAPA Probe Fast qPCR kit, Bioline). Primers and probe used for the detection of *P. melissococcoides* were designed for this study (Table S7) and we used primers and probe from Dainat et al. (2018) for *M. plutonius* detection (Table S7). The DNA amplification was performed with a BioRad CFx96 Real Time System. The cycling program was as follows: initial denaturation step at 95°C for 3 min, followed by 55 cycles of 3 s at 95°C and 30 s at 60°C. Samples were run in duplicate and their C_q_ value averaged. Samples were considered as positive when averaged C_q_ ≤ 55, and negative when no signal (C_q_ = N/A, no answer) was detected at the end of the amplification cycles.

Idiopathic samples (N=4) were collected from colonies displaying EFB symptoms, but in which *M. plutonius* and *P. larvae* (in two cases) were not detected by PCR and microscopy (Table S7).

### *P. melissococcoides* screening in the NCBI GenBank^®^ repository

The complete 16S rDNA sequence of *P. melissococcoides* 2.1 (GCA_944800085) was compared with these of *Paenibacillus* spp. (taxid 44249) available on GenBank^®^ (DDBJ/ENA/GenBank, accessed on 12.12.2022) using the BlastN tool (Zhang et al., 2000; Morgulis et al., 2008).

### Statistical analysis

Survival differences between experimental groups were illustrated with Kaplan-Meier survival curves and 95% confidence intervals. Differences in brood survival between the groups were tested using log-rank tests for two groups comparisons (bacterial inoculation versus control) and pairwise log-rank tests corrected by Bonferroni-Holm method for multiple comparisons (bacterial inoculation at different bacterial doses or larval ages versus controls). The significance level α was fixed at 0.05. The software R, version 4.1.0 was used for statistical tests. Kaplan-Meier survival curves were viewed using the “ggplot2” package and safe colourblind palette (Team, 2016; Wickham, 2016; Wong, 2011).

## Acknowledgments

We are grateful to Valérie Chaignat from the veterinary institute of Galli-Valerio, Lausanne, Etat de Vaud and to Daniela Grossar for providing us with samples for screening. We are also grateful to Walter Gasser, Bern cantonal bee inspector and his team, who supported us in the communication with the beekeepers during sample collections. We acknowledge Daniel Marzohl for the MALDI-TOF analyses and the support provided during data interpretation, Hiromi Imamoto for conducting some of the qPCR analyses and to Noah Squaratti for bacterial cultures and spore counts.

CRediT

Florine Ory: Conceptualization, Investigation, Formal Analysis, Data Curation, Writing - Original Draft, Writing - Review & Editing, Visualisation, Project administration.

Vincent Dietemann: Conceptualization, Writing - Original Draft, Writing - Review & Editing, Visualisation, Supervision, Project administration.

Alexandra Roetschi: Methodology, Investigation, Resources, Validation, Review & Editing.

Fabian Wenger: Methodology, Investigation, Data Curation, Validation, Review & Editing.

Oliver Würgler: Methodology, Investigation, Data Curation, Validation, Review & Editing.

Lauriane Braillard: Methodology, Investigation, Resources, Validation.

Jean-Daniel Charrière: Conceptualization, Writing - Review & Editing.

Benjamin Dainat: Conceptualization, Investigation, Validation, Writing - Original Draft, Writing - Review & Editing, Visualisation, Supervision, Project administration.

---

**Figure S1:**
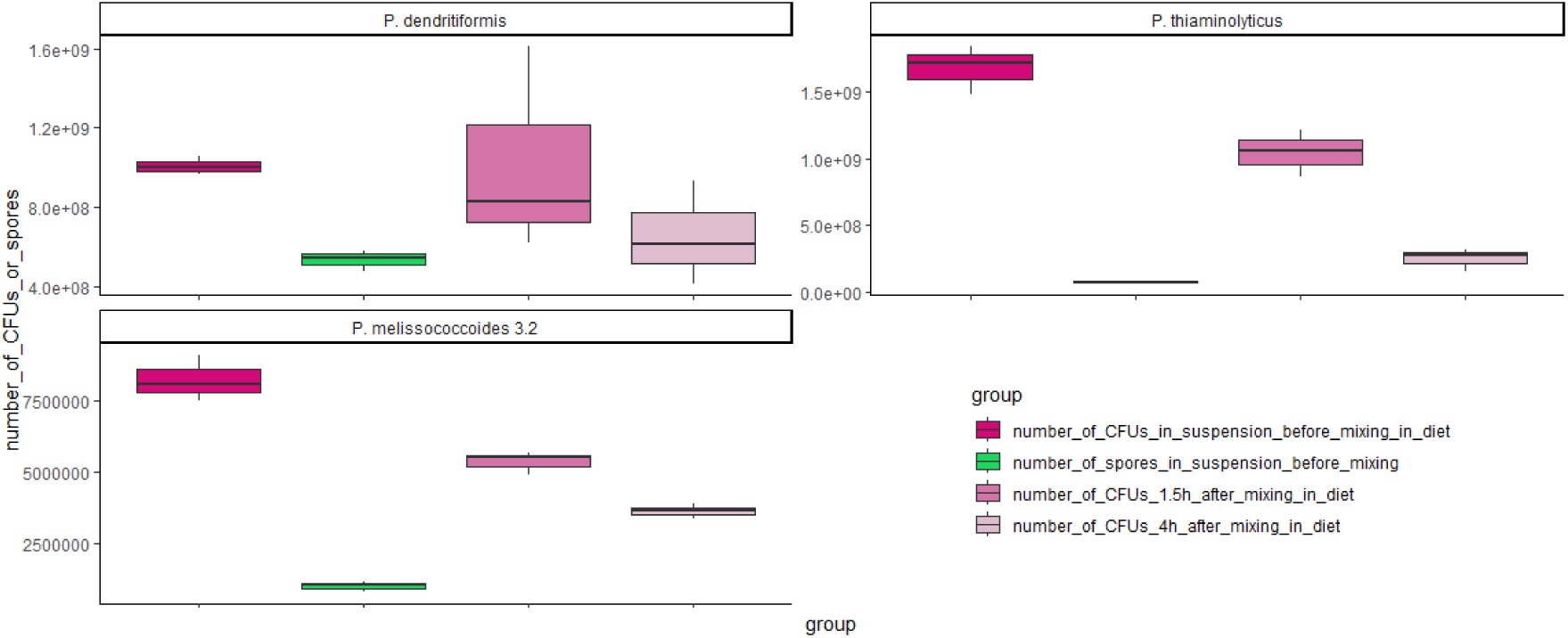
Number of colony forming units in suspensions of *Paenibacillus melissococcoides*, *Paenibacillus dendritiformis* and *Paenibacillus thiaminolyticus* immediately before, 1.5 and 4h after mixing with the diet. The number of spores in the suspensions was calculated from the percentage of spores evaluated by microscopy before mixing with the diet.

