## Supplementary Material for "Ecology and pathogenicity for honey bee brood of recently described Paenibacillus melissococcoides and comparison with Paenibacillus dendritiformis, Paenibacillus thiaminolyticus"

**Table S1.** Quantification by qPCR (copies/reaction) of *Paenibacillus melissococcoides* from pools of adult worker and larval. Samples were collected in 2021 from ten colonies (numbered 1 to 10) with or without European Foulbrood symptoms at the apiary at which the bacterium was first isolated in 2020.

| **Sample** | **Copies/reaction** | **Sample** | **Copies/reaction** | **EFB symptoms** |
| --- | --- | --- | --- | --- |
| adult workers 1 | 0.0E+00 | larvae 1 | 0.0E+00 | no |
| adult workers 2 | 2.6E+03 | larvae 2 | 2.3E+04 | yes |
| adult workers 3 | 1.7E+05 | larvae 3 | 2.1E+03 | yes |
| adult workers 4 | 9.8E+04 | larvae 4 | 1.1E+02 | yes |
| adult workers 5 | 1.6E+05 | larvae 5 | 1.0E+02 | yes |
| adult workers 6 | 3.5E+05 | larvae 6 | 5.8E+01 | yes |
| adult workers 7 | 5.8E+03 | larvae 7 | 1.5E+03 | yes |
| adult workers 8 | 5.1E+03 | larvae 8 | 4.0E+03 | yes |
| adult workers 9 | 4.9E+05 | larvae 9 | 3.4E+03 | yes |
| adult workers 10 | 4.8E+03 | larvae 10 | 1.0E+05 | yes |

**Table S2.** Log-Rank test results of the comparison of survival between a non-inoculated control and brood inoculated with three isolates of *Paenibacillus melissococcoides,* one isolate each of *Paenibacillus dendritiformis, Paenibacillus thiaminolyticus, Melissococcus plutonius and* reisolated *Paenibacillus melissococcoides 1.2.*

| **Brood treatment** | **N** | **Observed** | **Expected** | **(O-E)^2/E** | **(O-E)^2/V** | **df** | **p-value** |
| --- | --- | --- | --- | --- | --- | --- | --- |
| Control | 96 | 12 | 56 | 34.6 | 114 |  |  |
| *P. melissococcoides* 1.2 | 96 | 80 | 36 | 53.8 | 114 | 1 | <2e-16 |
| Control | 96 | 14 | 38.2 | 15.3 | 39 |  |  |
| *P. melissococcoides* 2.1 | 96 | 53 | 28.8 | 20.4 | 39 | 1 | 4e-10 |
| Control | 72 | 6 | 33.3 | 22.3 | 53.2 |  |  |
| *P. melissococcoides* 3.2 | 96 | 61 | 33.7 | 22.0 | 53.2 | 1 | 3e-13 |
| Control | 96 | 27 | 74.5 | 30.3 | 157 |  |  |
| *P. dendritiformis* LMG 21716 | 96 | 92 | 44.5 | 50.8 | 157 | 1 | <2e-16 |
| Control | 96 | 12 | 24.2 | 6.13 | 13.3 |  |  |
| *P. thiaminolyticus* DSM 7262 | 96 | 34 | 21.8 | 6.79 | 13.3 | 1 | 3e-04 |
| Control | 96 | 14 | 64.5 | 39.5 | 130 |  |  |
| *M. plutonius* CH 49.3 | 96 | 87 | 36.5 | 69.7 | 130 | 1 | <2e-16 |
| Control | 48 | 4 | 25.5 | 18.2 | 45.6 |  |  |
| *P. melissococcoides* 1.2 reisolated | 72 | 50 | 28.5 | 16.3 | 45.6 | 1 | 1e-11 |

**Table S3.** Expected and recovered number of colony forming units (CFUs) per larva estimated by plating the *Paenibacillus* spp. and *Melissococcus* *plutonius* inocula 1.5 h post-inoculation of one-, three- and five-days old larvae and of one-day old larvae with various bacterial doses. Plating quantified the maximal dose of viable bacteria to which the larvae were exposed. The ratio between expected and average recovered is also given.

| **Inoculum content** | **Larval age at inoculation (days)** | **Inoculum dose  (number of bacteria)** | **Mean ± SE number of CFUs recovered** | **Ratio** |
| --- | --- | --- | --- | --- |
| Control | 1 | 0 | 0 ± 0 | - |
| *M. plutonius* CH 49.3 | 1 | 2x10^5^ | 4.00x10^4^ ± 5470 | 5.0 |
| *P. thiaminolyticus* DSM 7262 | 1 | 2x10^5^ | 0 ± 0 | - |
| *P. dendritiformis* LMG 21716 | 1 | 2x10^5^ | 2.19x10^4^ ± 10100 | 9.1 |
| *P. melissococcoides* 1.2 | 1 | 2x10^5^ | 1.93x10^4^ ± 6500 | 10.4 |
| *P. melissococcoides* 2.1 | 1 | 2x10^5^ | 1.48x10^4^ ± 4370 | 13.5 |
| *P. melissococcoides* 3.2 | 1 | 2x10^5^ | 8.05x10^3^ ± 3210 | 24.8 |
| *P. melissococcoides* 3.2 | 1 | 2x10^4^ | 134 ± 1.3 | 149.3 |
| *P. melissococcoides* 3.2 | 1 | 2x10^3^ | 20.3 ± 0.2 | 100.0 |
| *P. melissococcoides* 3.2 | 3 | 2x10^5^ | 1.75x10^4^ ± 6120 | 11.4 |
| *P. melissococcoides* 3.2 | 5 | 2x10^5^ | 1.66x10^4^ ± 5600 | 12.0 |
| *P. melissococcoides* 1.2 reisolate | 1 | 2x10^5^ | 7.23x10^3^ ± 2670 | 27.7 |

**Table S4.** Concentration of *Paenibacillus thiaminolyticus* bacterial suspension estimated by plating before mixing with the diet to prepare the inoculum at the dose mentioned in Table S2.

| Inoculum content | Bacterial suspension concentration (number of CFUs ml^-1^) |
| --- | --- |
| *P. thiaminolyticus* DSM 7262 | 7.38x10^7^ ± 2.89x10^7^ |

**Table S5.** Average percentage (SD) of spores in stock suspensions of *Paenibacillus melissococcoides, Paenibacillus dendritiformis and Paenibacillus thyaminolyticus*, calculated based on hemocytometer counts.

| Bacteria species and strain | % spores |
| --- | --- |
| *Paenibacillus melissococoides* 3.2 | 12.0 (2.0) |
| *Paenibacillus dendritiformis* LMG 21726 | 53.0 (7.0) |
| *Paenibacillus thyaminolyticus* DSM 7262 | 4.4 (0.1) |

**Table S6.** Number of CFUs recovered from larvae (N=7) of three development stages experimentally infected with *P. melissococcoides* isolates 1.2 and 2.1.

|  | *P. melissococcoides* isolate | |
| --- | --- | --- |
| Larval stage | 1.2 | 2.1 |
| L3 | 660 | 420 |
| L4 #1 | 516 | 4572 |
| L4 #2 | 60 | - |
| L5 | 2.7x10^5^ | >2.2x10^4^ |

**Table S7. Screening results for *Paenibacillus melissococcoides* (2005-2021) in honey bee workers from *Melissococcus plutonius* negative and positive colonies.** Sample number, sampled apiary code, number of sampled / pooled colonies per apiary, sampling dates, colony sanitary status, Cq values and idiopathic cases are indicated. The mention 'positive' for *M. plutonius* designates samples previously identified as positive visually and by conventional PCR, but of which the DNA was too degraded for a qPCR in the framework of this study. When bacterium diagnostics was done by microscopy, it is indicated in the relevant column. By default, PCR results are given. Samples positive for *P. melissococcoides* are highlighted in yellow. Samples negative for *M. plutonius* are highlighted in orange. Idiopathic cases (negative for M. plutonius but showing symptoms) are highlighted in red in the last column. * indicates samples also microscopically negative for *Paenibacillus larvae*.

| **Sample** | **Apiary** | **Number of colonies** | **Sampling date** | **Colony status (visual determination)** | ***Paenibacillus melissococcoides* negative or positive (Cq mean)** | ***Melissococcus plutonius negative or positive* (Cq mean or microscopy)** |
| --- | --- | --- | --- | --- | --- | --- |
| 42 | SO7 | 1 | 2005 | healthy | negative | negative |
| 43 | BE130 | 1 | 2005 | healthy | negative | negative |
| 44 | BE120 | 1 | 2005 | healthy | negative | negative |
| 45 | SO8 | 1 | 2005 | healthy | negative | negative |
| 46 | SO12 | 1 | 2005 | healthy | negative | negative |
| 37 | BE3 | 1 | 2006 | healthy | negative | negative |
| 38 | VS1 | 1 | 2006 | healthy | negative | negative |
| 39 | BE121 | 1 | 2006 | healthy | negative | negative |
| 40 | BE122 | 1 | 2006 | healthy | negative | negative |
| 41 | GL1 | 1 | 2006 | healthy | negative | negative |
| 69 | SO0307 | 1 | 2007 | healthy | negative | negative |
| 80 | SO0907 | 1 | 2007 | healthy | negative | negative |
| 33 | BE0307 | 1 | 2007 | healthy | negative | negative |
| 34 | BE0407 | 1 | 2007 | healthy | negative | negative |
| 35 | BE0507 | 1 | 2007 | healthy | negative | negative |
| 36 | SO0906 | 1 | 2007 | healthy | negative | negative |
| 107 | BE21108 | 1 | 2008 | healthy | negative | negative |
| 27 | GSR1 | 1 | 2008 | healthy | negative | negative |
| 28 | BE20508 | 1 | 2008 | healthy | negative | negative |
| 29 | EW1 | 15 | 2008 | healthy | negative | negative |
| 30 | SB1 | 16 | 2008 | healthy | negative | negative |
| 31 | GT1 | 8 | 2008 | healthy | negative | negative |
| 32 | BSR1 | 6 | 2008 | healthy | negative | negative |
| 20 | RMO1 | 2 | 2009 | healthy | negative | negative |
| 21 | EM1 | 5 | 2009 | healthy | negative | negative |
| 22 | VF | 3 | 2009 | healthy | negative | negative |
| 23 | TW | 1 | 2009 | healthy | negative | negative |
| 24 | VC | 1 | 2009 | healthy | negative | negative |
| 25 | ZK | 2 | 2009 | healthy | negative | negative |
| 26 | LW1 | 3 | 2009 | healthy | negative | negative |
| 15 | HB1 | 2 | 2010 | healthy | negative | negative |
| 16 | HB2 | 3 | 2010 | healthy | negative | negative |
| 17 | RF1 | 6 | 2010 | healthy | negative | negative |
| 18 | EK1 | 1 | 2010 | healthy | negative | negative |
| 19 | ODJT1 | 32 | 2010 | healthy | negative | negative |
| 10 | WW1 | 4 | 2011 | healthy | negative | negative |
| 11 | BCJ | 5 | 2011 | healthy | negative | negative |
| 12 | CB1 | 3 | 2011 | healthy | negative | negative |
| 13 | BC1 | 3 | 2011 | healthy | negative | negative |
| 14 | BC2 | 3 | 2011 | healthy | negative | negative |
| 6 | LF6 | 1 | 2019 | healthy | negative | negative |
| 7 | LW2 | 1 | 2019 | healthy | negative | negative |
| 8 | LW3 | 1 | 2019 | healthy | negative | negative |
| 9 | LW4 | 1 | 2019 | healthy | negative | negative |
| 1 | LF1 | 1 | 2020 | healthy | negative | negative |
| 2 | LF2 | 1 | 2020 | healthy | negative | negative |
| 3 | LF3 | 1 | 2020 | healthy | negative | negative |
| 4 | LF4 | 1 | 2020 | healthy | negative | negative |
| 5 | LF5 | 1 | 2020 | healthy | negative | negative |
| 1 | VD1272 | 1 | 2021 | healthy | negative | negative |
| 4 | VD1406 | 1 | 2021 | EFB symptomatic | negative | negative |
| 70 | VD34421 | 1 | 2021 | EFB symptomatic | negative | negative |
| 1 | BE1 | 1 | 2005 | EFB symptomatic | negative | 22.00 |
| 2 | BE1 | 1 | 2005 | EFB symptomatic | negative | 16.31 |
| 3 | BE1 | 1 | 2005 | EFB symptomatic | negative | 17.43 |
| 4 | BE1 | 1 | 2005 | EFB symptomatic | negative | 15.33 |
| 5 | BE2 | 1 | 2005 | EFB symptomatic | negative | 16.21 |
| 6 | BE2 | 1 | 2005 | EFB symptomatic | negative | 16.69 |
| 7 | BE2 | 1 | 2005 | EFB symptomatic | negative | 15.56 |
| 8 | BE3 | 1 | 2005 | EFB symptomatic | negative | 14.88 |
| 9 | BE3 | 1 | 2005 | EFB symptomatic | negative | 18.60 |
| 10 | BE3 | 1 | 2005 | EFB symptomatic | negative | 16.43 |
| 11 | BE120 | 1 | 2005 | EFB symptomatic | negative | 18.49 |
| 12 | BE120 | 1 | 2005 | EFB symptomatic | negative | 14.50 |
| 13 | BE120 | 1 | 2005 | EFB symptomatic | negative | 15.39 |
| 15 | BE121 | 1 | 2005 | EFB symptomatic | negative | 16.01 |
| 16 | BE121 | 1 | 2005 | EFB symptomatic | negative | 15.42 |
| 17 | BE121 | 1 | 2005 | EFB symptomatic | negative | 14.88 |
| 18 | BE122 | 1 | 2005 | EFB symptomatic | negative | 16.26 |
| 19 | BE122 | 1 | 2005 | EFB symptomatic | negative | 13.71 |
| 20 | BE122 | 1 | 2005 | EFB symptomatic | negative | 17.22 |
| 21 | BE123 | 1 | 2005 | EFB symptomatic | negative | 19.23 |
| 22 | BE123 | 1 | 2005 | EFB symptomatic | negative | 16.71 |
| 23 | BE123 | 1 | 2005 | EFB symptomatic | negative | 17.35 |
| 24 | BE124 | 1 | 2005 | EFB symptomatic | negative | 16.04 |
| 25 | BE124 | 1 | 2005 | EFB symptomatic | negative | 16.58 |
| 26 | BE134 | 1 | 2005 | EFB symptomatic | negative | 20.31 |
| 27 | BE134 | 1 | 2005 | EFB symptomatic | negative | 20.52 |
| 28 | SO7 | 1 | 2005 | EFB symptomatic | negative | 22.09 |
| 29 | SO7 | 1 | 2005 | EFB symptomatic | negative | 14.02 |
| 30 | SO7 | 1 | 2005 | EFB symptomatic | negative | 14.32 |
| 31 | SO8 | 1 | 2005 | EFB symptomatic | negative | 16.64 |
| 32 | SO8 | 1 | 2005 | EFB symptomatic | negative | 18.89 |
| 33 | SO8 | 1 | 2005 | EFB symptomatic | negative | 15.41 |
| 34 | SO12 | 1 | 2005 | EFB symptomatic | negative | 16.91 |
| 35 | SO12 | 1 | 2005 | EFB symptomatic | negative | 14.70 |
| 36 | VS2 | 1 | 2006 | EFB symptomatic | negative | 21.23 |
| 37 | VS2 | 1 | 2006 | EFB symptomatic | negative | 23.60 |
| 38 | VS2 | 1 | 2006 | EFB symptomatic | negative | 22.60 |
| 39 | GL1 | 1 | 2006 | EFB symptomatic | negative | 22.23 |
| 40 | SO6 | 1 | 2006 | EFB symptomatic | negative | 21.77 |
| 187 | SO09061 | 1 | 2006 | EFB symptomatic | negative | 16.29 |
| 188 | SO09062 | 1 | 2006 | EFB symptomatic | negative | 16.65 |
| 189 | SO09063 | 1 | 2006 | EFB symptomatic | negative | 20.44 |
| 41 | BE01071 | 1 | 2007 | EFB symptomatic | negative | 21.98 |
| 42 | BE01072 | 1 | 2007 | EFB symptomatic | negative | 27.64 |
| 43 | BE01073 | 1 | 2007 | EFB symptomatic | negative | 26.00 |
| 44 | BE02071 | 1 | 2007 | EFB symptomatic | negative | 22.07 |
| 45 | BE02072 | 1 | 2007 | EFB symptomatic | negative | 28.55 |
| 46 | BE02073 | 1 | 2007 | EFB symptomatic | negative | 28.10 |
| 47 | BE03071 | 1 | 2007 | EFB symptomatic | negative | 18.65 |
| 48 | BE06071 | 1 | 2007 | EFB symptomatic | negative | 19.06 |
| 49 | BE06072 | 1 | 2007 | EFB symptomatic | negative | 23.11 |
| 50 | BE06073 | 1 | 2007 | EFB symptomatic | negative | 22.28 |
| 51 | BE06074 | 1 | 2007 | EFB symptomatic | negative | 20.42 |
| 52 | BE06075 | 1 | 2007 | EFB symptomatic | negative | 17.29 |
| 53 | BE07071 | 1 | 2007 | EFB symptomatic | negative | 18.42 |
| 54 | BE07072 | 1 | 2007 | EFB symptomatic | negative | 21.20 |
| 55 | BE07073 | 1 | 2007 | EFB symptomatic | negative | 24.76 |
| 56 | BE08071 | 1 | 2007 | EFB symptomatic | negative | 20.97 |
| 57 | BE08072 | 1 | 2007 | EFB symptomatic | negative | 29.81 |
| 58 | BE09071 | 1 | 2007 | EFB symptomatic | negative | 29.72 |
| 59 | BE09072 | 1 | 2007 | EFB symptomatic | negative | 22.01 |
| 60 | SO02071 | 1 | 2007 | EFB symptomatic | negative | 20.55 |
| 61 | SO02072 | 1 | 2007 | EFB symptomatic | negative | 22.51 |
| 62 | SO02073 | 1 | 2007 | EFB symptomatic | negative | 21.54 |
| 63 | SO02074 | 1 | 2007 | EFB symptomatic | negative | 21.77 |
| 64 | SO02075 | 1 | 2007 | EFB symptomatic | negative | 23.11 |
| 65 | SO02076 | 1 | 2007 | EFB symptomatic | negative | 18.73 |
| 66 | SO3071 | 1 | 2007 | EFB symptomatic | negative | 21.33 |
| 67 | SO3072 | 1 | 2007 | EFB symptomatic | negative | 28.61 |
| 68 | SO3073 | 1 | 2007 | EFB symptomatic | negative | 37.12 |
| 70 | SO4071 | 1 | 2007 | EFB symptomatic | negative | 21.02 |
| 71 | SO4072 | 1 | 2007 | EFB symptomatic | negative | 20.61 |
| 72 | SO4073 | 1 | 2007 | EFB symptomatic | negative | 18.68 |
| 73 | SO4074 | 1 | 2007 | EFB symptomatic | negative | 21.99 |
| 74 | SO5071 | 1 | 2007 | EFB symptomatic | negative | 23.30 |
| 75 | SO5072 | 1 | 2007 | EFB symptomatic | negative | 20.41 |
| 76 | SO5073 | 1 | 2007 | EFB symptomatic | negative | 22.44 |
| 77 | SO6071 | 1 | 2007 | EFB symptomatic | negative | 24.09 |
| 78 | SO6072 | 1 | 2007 | EFB symptomatic | negative | 23.87 |
| 79 | SO6073 | 1 | 2007 | EFB symptomatic | negative | 20.86 |
| 81 | SO9071 | 1 | 2007 | EFB symptomatic | negative | 20.66 |
| 82 | SO9072 | 1 | 2007 | EFB symptomatic | negative | 14.97 |
| 83 | SO9073 | 1 | 2007 | EFB symptomatic | negative | 16.41 |
| 181 | BE00071 | 1 | 2007 | EFB symptomatic | negative | 15.46 |
| 182 | BE00072 | 1 | 2007 | EFB symptomatic | negative | 33.19 |
| 183 | BE00073 | 1 | 2007 | EFB symptomatic | negative | 15.96 |
| 185 | BE01071 | 1 | 2007 | EFB symptomatic | negative | 16.13 |
| 186 | BE01072 | 1 | 2007 | EFB symptomatic | negative | 17.26 |
| 190 | NA1 | 1 | 2008 | EFB symptomatic | negative | 34.42 |
| 191 | NA2 | 1 | 2008 | EFB symptomatic | negative | 31.12 |
| 184 | BE00073 | 1 | 2008 | EFB symptomatic | negative | 19.80 |
| 84 | G 396 | 1 | 2008 | EFB symptomatic | negative | 14.24 |
| 85 | G 396 | 1 | 2008 | EFB symptomatic | negative | 17.74 |
| 86 | G 396 | 1 | 2008 | EFB symptomatic | negative | 26.48 |
| 87 | G 372 | 1 | 2008 | EFB symptomatic | negative | 16.24 |
| 88 | G 372 | 1 | 2008 | EFB symptomatic | negative | 17.57 |
| 89 | G 375 | 1 | 2008 | EFB symptomatic | negative | 33.42 |
| 90 | G 375 | 1 | 2008 | EFB symptomatic | negative | 20.19 |
| 91 | G 375 | 1 | 2008 | EFB symptomatic | negative | 14.57 |
| 92 | G 395 | 1 | 2008 | EFB symptomatic | negative | 12.79 |
| 93 | G 395 | 1 | 2008 | EFB symptomatic | negative | 15.33 |
| 94 | G 395 | 1 | 2008 | EFB symptomatic | negative | 14.88 |
| 95 | G 395 | 1 | 2008 | EFB symptomatic | negative | 13.80 |
| 96 | BE201081 | 1 | 2008 | EFB symptomatic | negative | 17.85 |
| 97 | BE201082 | 1 | 2008 | EFB symptomatic | negative | 15.12 |
| 98 | BE201083 | 1 | 2008 | EFB symptomatic | negative | 22.48 |
| 99 | BE202081 | 1 | 2008 | EFB symptomatic | negative | 34.70 |
| 100 | BE202082 | 1 | 2008 | EFB symptomatic | negative | 17.16 |
| 101 | BE202083 | 1 | 2008 | EFB symptomatic | negative | 16.24 |
| 102 | BE202084 | 1 | 2008 | EFB symptomatic | negative | 16.93 |
| 103 | BE203081 | 1 | 2008 | EFB symptomatic | negative | 23.49 |
| 104 | BE203082 | 1 | 2008 | EFB symptomatic | negative | 19.44 |
| 105 | BE203083 | 1 | 2008 | EFB symptomatic | negative | 23.66 |
| 106 | BE204081 | 1 | 2008 | EFB symptomatic | negative | 16.89 |
| 108 | BE211081 | 1 | 2008 | EFB symptomatic | negative | 18.92 |
| 109 | BE211082 | 1 | 2008 | EFB symptomatic | negative | 17.96 |
| 110 | BE211083 | 1 | 2008 | EFB symptomatic | negative | 17.26 |
| 111 | BE212081 | 1 | 2008 | EFB symptomatic | negative | 27.36 |
| 112 | BE213081 | 1 | 2008 | EFB symptomatic | negative | 16.89 |
| 113 | BE213082 | 3 | 2008 | EFB symptomatic | negative | 19.62 |
| 114 | BE214081 | 1 | 2008 | EFB symptomatic | negative | 18.82 |
| 115 | BE217081 | 1 | 2008 | EFB symptomatic | negative | 19.62 |
| 116 | BE217082 | 1 | 2008 | EFB symptomatic | negative | 17.50 |
| 117 | BE217083 | 1 | 2008 | EFB symptomatic | negative | 35.04 |
| 118 | BE360081 | 1 | 2008 | EFB symptomatic | negative | 19.96 |
| 119 | BE360082 | 1 | 2008 | EFB symptomatic | negative | 27.06 |
| 120 | SO1081 | 1 | 2008 | EFB symptomatic | negative | 17.08 |
| 121 | SO1082 | 1 | 2008 | EFB symptomatic | negative | 20.23 |
| 122 | SO1083 | 1 | 2008 | EFB symptomatic | negative | 21.42 |
| 123 | SO2081 | 1 | 2008 | EFB symptomatic | negative | 24.73 |
| 124 | SO3081 | 1 | 2008 | EFB symptomatic | negative | 19.14 |
| 125 | SO3082 | 1 | 2008 | EFB symptomatic | negative | 25.96 |
| 126 | SO4081 | 1 | 2008 | EFB symptomatic | negative | 18.17 |
| 127 | SO4082 | 1 | 2008 | EFB symptomatic | negative | 20.16 |
| 128 | SO5081 | 1 | 2008 | EFB symptomatic | negative | 15.18 |
| 129 | SO5082 | 1 | 2008 | EFB symptomatic | negative | 24.50 |
| 130 | SO5083 | 1 | 2008 | EFB symptomatic | negative | 23.85 |
| 131 | SO7081 | 1 | 2008 | EFB symptomatic | negative | 16.46 |
| 132 | SO7082 | 1 | 2008 | EFB symptomatic | negative | 20.25 |
| 142 | MS1 | 6 | 2008 | EFB symptomatic | negative | 22.48 |
| 143 | SB2 | 16 | 2008 | EFB symptomatic | negative | 28.21 |
| 144 | BA1 | 11 | 2008 | EFB symptomatic | negative | 18.10 |
| 145 | GS1 | 5 | 2008 | EFB symptomatic | negative | 20.59 |
| 146 | HGL1 | 15 | 2008 | EFB symptomatic | negative | 23.91 |
| 147 | IS1 | 41 | 2008 | EFB symptomatic | negative | 19.23 |
| 148 | KL1 | 7 | 2008 | EFB symptomatic | negative | 20.33 |
| 149 | VW1 | 12 | 2008 | EFB symptomatic | negative | 17.53 |
| 165 | TH1 | 9 | 2008 | EFB symptomatic | negative | 21.96 |
| 166 | LB1 | 9 | 2008 | EFB symptomatic | negative | 20.43 |
| 167 | BM1 | 5 | 2008 | EFB symptomatic | negative | 17.97 |
| 150 | WM1 | 14 | 2009 | EFB symptomatic | negative | 20.45 |
| 151 | WD1 | 4 | 2009 | EFB symptomatic | negative | 23.99 |
| 152 | ANG1 | 22 | 2009 | EFB symptomatic | negative | 19.44 |
| 153 | HB2 | 11 | 2009 | EFB symptomatic | negative | 18.17 |
| 154 | DRF1 | 12 | 2009 | EFB symptomatic | negative | 16.88 |
| 155 | TM1 | 15 | 2009 | EFB symptomatic | negative | 21.81 |
| 156 | LM1 | 16 | 2009 | EFB symptomatic | negative | 20.96 |
| 157 | EOO1 | 22 | 2009 | EFB symptomatic | negative | 17.47 |
| 158 | LHRBST | 22 | 2009 | EFB symptomatic | negative | 19.24 |
| 159 | AD1 | 17 | 2009 | EFB symptomatic | negative | 22.15 |
| 160 | BRT | 6 | 2009 | EFB symptomatic | negative | 19.68 |
| 161 | GRSU | 10 | 2009 | EFB symptomatic | negative | 23.52 |
| 162 | RAAF | 6 | 2009 | EFB symptomatic | negative | 22.28 |
| 163 | KPG | 18 | 2009 | EFB symptomatic | negative | 18.01 |
| 164 | BLGOH | 14 | 2009 | EFB symptomatic | negative | 24.10 |
| 168 | NA | 7 | 2009 | EFB symptomatic | negative | 20.40 |
| 169 | GGL | 11 | 2009 | EFB symptomatic | negative | 17.69 |
| 170 | GUGL | 6 | 2009 | EFB symptomatic | negative | 31.30 |
| 171 | LBSTRB | 3 | 2009 | EFB symptomatic | negative | 17.38 |
| 172 | KRK1 | 7 | 2009 | EFB symptomatic | negative | 18.14 |
| 173 | KK2 | 10 | 2009 | EFB symptomatic | negative | 20.85 |
| 174 | KRH1 | 3 | 2009 | EFB symptomatic | negative | 23.58 |
| 175 | HISA | 10 | 2009 | EFB symptomatic | negative | 32.05 |
| 176 | AESA | 12 | 2009 | EFB symptomatic | negative | 19.01 |
| 177 | SCWRF | 6 | 2009 | EFB symptomatic | negative | 32.57 |
| 179 | VC2 | 14 | 2009 | EFB symptomatic | negative | 25.63 |
| 180 | VC3 | 1 | 2009 | EFB symptomatic | negative | 25.59 |
| 192 | NA3 | 1 | 2009 | EFB symptomatic | negative | 16.48 |
| 193 | NA4 | 1 | 2009 | EFB symptomatic | negative | 19.28 |
| 194 | NA5 | 1 | 2009 | EFB symptomatic | negative | 18.67 |
| 195 | NA6 | 1 | 2009 | EFB symptomatic | negative | 17.35 |
| 196 | NA7 | 1 | 2009 | EFB symptomatic | negative | 16.69 |
| 197 | NA8 | 1 | 2009 | EFB symptomatic | negative | 19.81 |
| 198 | NA9 | 1 | 2009 | EFB symptomatic | negative | 17.67 |
| 199 | NA10 | 1 | 2009 | EFB symptomatic | negative | 16.82 |
| 200 | RF1 | 1 | 2010 | EFB symptomatic | negative | 18.49 |
| 201 | RF2 | 1 | 2010 | EFB symptomatic | negative | 18.71 |
| 202 | RF3 | 1 | 2010 | EFB symptomatic | negative | 17.50 |
| 203 | SF1 | 1 | 2010 | EFB symptomatic | negative | 18.92 |
| 204 | RF4 | 1 | 2010 | EFB symptomatic | negative | 18.57 |
| 205 | RF5 | 1 | 2010 | EFB symptomatic | negative | 15.07 |
| 178 | SF1 | 8 | 2010 | EFB symptomatic | negative | 18.06 |
| 133 | BC21 | 3 | 2010 | EFB symptomatic | negative | 23.42 |
| 134 | BC22 | 3 | 2010 | EFB symptomatic | negative | 28.41 |
| 135 | BC23 | 4 | 2010 | EFB symptomatic | negative | 19.09 |
| 136 | BC24 | 4 | 2010 | EFB symptomatic | negative | 25.88 |
| 137 | BC25 | 1 | 2010 | EFB symptomatic | negative | 18.38 |
| 138 | BC26 | 1 | 2010 | EFB symptomatic | negative | 20.06 |
| 139 | BC27 | 1 | 2010 | EFB symptomatic | negative | 17.82 |
| 140 | BC28 | 1 | 2010 | EFB symptomatic | negative | 19.70 |
| 141 | BC29 | 1 | 2010 | EFB symptomatic | negative | 17.55 |
| 120 | 508-002 | 1 | 2013 | EFB symptomatic | negative | positive |
| 121 | 07.04.2169 | 1 | 2013 | EFB symptomatic | negative | 15.975 |
| 122 | 709-005 | 1 | 2013 | EFB symptomatic | negative | 16.735 |
| 123 | 710-002 | 1 | 2013 | EFB symptomatic | negative | 15.235 |
| 124 | No4_2013 | 2 | 2013 | EFB symptomatic | negative | 10.22 |
| 125 | No3_2013 | 1 | 2013 | EFB symptomatic | negative | 6.8875 |
| 126 | 514-006 | 1 | 2013 | EFB symptomatic | negative | positive |
| 127 | 731-006 | 1 | 2013 | EFB symptomatic | negative | positive |
| 128 | 709-010 | 1 | 2013 | EFB symptomatic | negative | 14.29 |
| 129 | 514-007 | 1 | 2013 | EFB symptomatic | negative | 16.045 |
| 130 | 528-002 | 1 | 2013 | EFB symptomatic | negative | 18.64 |
| 131 | 530-002 | 1 | 2013 | EFB symptomatic | negative | 10.54 |
| 132 | 723-007 | 1 | 2013 | EFB symptomatic | negative | 17.16 |
| 133 | 710-003 | 1 | 2013 | EFB symptomatic | negative | 13.865 |
| 134 | 618-007 | 1 | 2013 | EFB symptomatic | negative | 8.98 |
| 135 | 607-002 | 1 | 2013 | EFB symptomatic | negative | 10.68 |
| 136 | 620-002 | 2 | 2013 | EFB symptomatic | negative | positive |
| 137 | 730-003 | 1 | 2013 | EFB symptomatic | negative | 19.725 |
| 138 | 808-002 | 1 | 2013 | EFB symptomatic | negative | 18.245 |
| 139 | 611-009 | 1 | 2013 | EFB symptomatic | negative | 17.155 |
| 140 | 424-003 | 1 | 2013 | EFB symptomatic | negative | 14.455 |
| 141 | 703-006 | 1 | 2013 | EFB symptomatic | negative | 8.49 |
| 142 | 819-008 | 1 | 2013 | EFB symptomatic | negative | 8.685 |
| 143 | 830-001 | 1 | 2013 | EFB symptomatic | negative | positive |
| 144 | 611-011 | 1 | 2013 | EFB symptomatic | 30.69 | 15.8125 |
| 145 | 514-005 | 1 | 2013 | EFB symptomatic | negative | 8.285 |
| 146 | 503-004 | 1 | 2013 | EFB symptomatic | negative | 7.6975 |
| 147 | No1_2013 | 1 | 2013 | EFB symptomatic | negative | 20.1 |
| 148 | 503-003 | 1 | 2013 | EFB symptomatic | negative | positive |
| 149 | 514-003 | 1 | 2013 | EFB symptomatic | negative | positive |
| 150 | 710-004 | 1 | 2013 | EFB symptomatic | negative | positive |
| 151 | 730-004 | 1 | 2013 | EFB symptomatic | negative | 19.145 |
| 152 | 529-006 | 1 | 2013 | EFB symptomatic | negative | positive |
| 153 | 425-006 | 1 | 2013 | EFB symptomatic | negative | 8.035 |
| 154 | 813-030 | 1 | 2013 | EFB symptomatic | negative | 8.555 |
| 155 | 903-007 | 1 | 2013 | EFB symptomatic | negative | 9.525 |
| 156 | 716-001 | 1 | 2013 | EFB symptomatic | negative | 17.07 |
| 157 | 507-004 | 1 | 2013 | EFB symptomatic | negative | 17.075 |
| 158 | 516-005 | 1 | 2013 | EFB symptomatic | negative | positive |
| 159 | 07.16.2179 | 1 | 2013 | EFB symptomatic | negative | 9.45 |
| 160 | 614-001 | 1 | 2013 | EFB symptomatic | negative | 10.26 |
| 161 | 611-010 | 1 | 2013 | EFB symptomatic | negative | 8.255 |
| 162 | 611-012 | 1 | 2013 | EFB symptomatic | negative | 20.67 |
| 163 | 529-004 | 1 | 2013 | EFB symptomatic | negative | positive |
| 164 | 702-010 | 1 | 2013 | EFB symptomatic | negative | positive |
| 165 | 626-003 | 1 | 2013 | EFB symptomatic | negative | 12.27 |
| 166 | 07.11.2173 | 1 | 2013 | EFB symptomatic | negative | 11.02 |
| 167 | 712-002 | 1 | 2013 | EFB symptomatic | negative | 14.96 |
| 168 | 610-009 | 1 | 2013 | EFB symptomatic | negative | 6.945 |
| 169 | 611-013 | 1 | 2013 | EFB symptomatic | negative | positive |
| 170 | 514-004 | 1 | 2013 | EFB symptomatic | negative | 18.885 |
| 171 | 703-003 | 1 | 2013 | EFB symptomatic | negative | 15.85 |
| 172 | KSSM | 1 | 2013 | EFB symptomatic | negative | 8.005 |
| 173 | 430-002 | 2 | 2013 | EFB symptomatic | negative | 8.23 |
| 174 | 619-001 | 1 | 2013 | EFB symptomatic | negative | 12.1 |
| 175 | 710-006 | 1 | 2013 | EFB symptomatic | negative | 12.615 |
| 176 | 507-003 | 1 | 2013 | EFB symptomatic | negative | positive |
| 177 | 614-002 | 1 | 2013 | EFB symptomatic | negative | 13.12 |
| 178 | 514-008 | 1 | 2013 | EFB symptomatic | negative | positive |
| 179 | 625-003 | 2 | 2013 | EFB symptomatic | negative | 9.93 |
| 180 | 625-004 | 1 | 2013 | EFB symptomatic | negative | 8.505 |
| 181 | 514-002 | 2 | 2013 | EFB symptomatic | negative | positive |
| 182 | 723-002 | 2 | 2013 | EFB symptomatic | negative | positive |
| 183 | 730-001 | 1 | 2013 | EFB symptomatic | negative | positive |
| 184 | 516-006 | 1 | 2013 | EFB symptomatic | negative | positive |
| 185 | 503-001 | 1 | 2013 | EFB symptomatic | negative | 7.455 |
| 186 | 611-014 | 1 | 2013 | EFB symptomatic | negative | positive |
| 187 | FLM No.1 | 1 | 2013 | EFB symptomatic | negative | positive |
| 188 | 613-002 | 1 | 2013 | EFB symptomatic | negative | 15.74 |
| 189 | 516-007 | 1 | 2013 | EFB symptomatic | negative | 20.97 |
| 190 | 516-008 | 1 | 2013 | EFB symptomatic | negative | 10.605 |
| 191 | 724-004 | 1 | 2013 | EFB symptomatic | negative | 15.17 |
| 192 | 716-003 | 1 | 2013 | EFB symptomatic | negative | 14.08 |
| 193 | 710-005 | 1 | 2013 | EFB symptomatic | negative | 14.57 |
| 194 | 516-010 | 1 | 2013 | EFB symptomatic | negative | positive |
| 195 | 620-003 | 1 | 2013 | EFB symptomatic | negative | 14.33 |
| 196 | 716-002 | 1 | 2013 | EFB symptomatic | negative | 18.215 |
| 197 | 704-003 | 1 | 2013 | EFB symptomatic | negative | 11.075 |
| IFB 1 | DV33952.2 | 1 | 2014 | EFB symptomatic | negative | 36.3 |
| EFB 1 | DV33951 | 1 | 2014 | EFB symptomatic | negative | 36.5 |
| EFB 3 | DV33955.1 | 1 | 2014 | EFB symptomatic | 39.5 | 36.6 |
| EFB 4 | DV33816.1 | 1 | 2014 | EFB symptomatic | negative | 34.4 |
| EFB5 | YC | 1 | 2014 | EFB symptomatic | negative | 36 |
| IFB 3 | DV32449 | 1 | 2015 | EFB symptomatic | negative | negative |
| EFB 7 | DV32450.6 | 1 | 2015 | EFB symptomatic | negative | positive |
| IFB 2 | DV32316 | 1 | 2015 | EFB symptomatic | 39.2 | negative |
| EFB 6 | DV54377 | 1 | 2015 | EFB symptomatic | 40.3 | positive |
| IFB 10 | VD47928 | 1 | 2018 | EFB symptomatic | 38.4 | negative microscopy* |
| IFB 9 | VD28438 | 1 | 2020 | EFB symptomatic | negative | negative microscopy |
| EFB 10 | VD24255 | 1 | 2020 | EFB symptomatic | 38.4 | positive microscopy* |
| EFB 9 | VD21343 | 1 | 2020 | EFB symptomatic | 38.1 | positive microscopy* |
| 2 | VD13417 | 1 | 2021 | EFB symptomatic | negative | 34.4 |
| 3 | VD13767 | 1 | 2021 | EFB symptomatic | negative | 39.8 |
| 5 | VD14591 | 1 | 2021 | EFB symptomatic | negative | 20.4 |
| 6 | VD14589 | 1 | 2021 | EFB symptomatic | negative | 19 |
| 7 | VD15098 | 1 | 2021 | EFB symptomatic | negative | 27.1 |
| 8 | VD15099 | 1 | 2021 | EFB symptomatic | negative | 32.7 |
| 9 | VD15100 | 1 | 2021 | EFB symptomatic | negative | 20.6 |
| 10 | VD15826 | 1 | 2021 | EFB symptomatic | negative | 39.5 |
| 11 | VD16220 | 1 | 2021 | EFB symptomatic | negative | 38 |
| 12 | VD16526 | 1 | 2021 | EFB symptomatic | negative | 18.5 |
| 13 | VD16525/1 | 1 | 2021 | EFB symptomatic | negative | 36.7 |
| 14 | VD16525/2 | 1 | 2021 | EFB symptomatic | negative | 36.6 |
| 15 | VD16527 | 1 | 2021 | EFB symptomatic | negative | 19.4 |
| 16 | VD16749 | 1 | 2021 | EFB symptomatic | 30.9 | 18.4 |
| 17 | VD16772/1 | 1 | 2021 | EFB symptomatic | negative | 18.6 |
| 18 | VD17490 | 1 | 2021 | EFB symptomatic | negative | 34.5 |
| 19 | VD17522/1 | 1 | 2021 | EFB symptomatic | negative | 27.7 |
| 20 | VD17672 | 1 | 2021 | EFB symptomatic | negative | 32.3 |
| 21 | VD17673 | 1 | 2021 | EFB symptomatic | negative | 23.1 |
| 22 | VD17884 | 1 | 2021 | EFB symptomatic | negative | 16.3 |
| 23 | VD18056 | 1 | 2021 | EFB symptomatic | negative | 21 |
| 24 | VD18331 | 1 | 2021 | EFB symptomatic | negative | 17.5 |
| 25 | VD18329/1 | 1 | 2021 | EFB symptomatic | negative | 22.3 |
| 26 | VD18325/1 | 1 | 2021 | EFB symptomatic | negative | 32.7 |
| 27 | VD18529 | 1 | 2021 | EFB symptomatic | negative | 18.7 |
| 28 | VD18651 | 1 | 2021 | EFB symptomatic | negative | 17.2 |
| 29 | VD18649/1 | 1 | 2021 | EFB symptomatic | negative | 33.1 |
| 30 | VD19817/1 | 1 | 2021 | EFB symptomatic | negative | 36.2 |
| 31 | VD19825 | 1 | 2021 | EFB symptomatic | negative | 22.2 |
| 32 | VD19818/1 | 1 | 2021 | EFB symptomatic | negative | 20.3 |
| 33 | VD20427 | 1 | 2021 | EFB symptomatic | negative | 20 |
| 34 | VD20697/1 | 1 | 2021 | EFB symptomatic | negative | 19.6 |
| 35 | VD20703 | 1 | 2021 | EFB symptomatic | negative | 23.3 |
| 36 | VD21025 | 1 | 2021 | EFB symptomatic | negative | 37.4 |
| 37 | VD21392 | 1 | 2021 | EFB symptomatic | negative | 42.8 |
| 38 | VD21393 | 1 | 2021 | EFB symptomatic | negative | 40.6 |
| 39 | VD21394 | 1 | 2021 | EFB symptomatic | negative | 34.5 |
| 40 | VD21625 | 1 | 2021 | EFB symptomatic | negative | 21.4 |
| 41 | VD21798 | 1 | 2021 | EFB symptomatic | negative | 37.9 |
| 42 | NA11 | 1 | 2021 | EFB symptomatic | negative | 17 |
| 43 | VD22428 | 1 | 2021 | EFB symptomatic | negative | 22.2 |
| 44 | VD22637 | 1 | 2021 | EFB symptomatic | negative | 16.4 |
| 45 | VD22636 | 1 | 2021 | EFB symptomatic | negative | 39.1 |
| 46 | VD22635-1 | 1 | 2021 | EFB symptomatic | 26.4 | 20.3 |
| 47 | VD22635-2 | 1 | 2021 | EFB symptomatic | negative | 34.8 |
| 48 | VD22635-2 | 1 | 2021 | EFB symptomatic | negative | 36 |
| 49 | VD23058 | 1 | 2021 | EFB symptomatic | negative | 28 |
| 50 | VD23061 | 1 | 2021 | EFB symptomatic | negative | 19.5 |
| 51 | VD23708 | 1 | 2021 | EFB symptomatic | negative | 20.5 |
| 52 | VD23885 | 1 | 2021 | EFB symptomatic | negative | 22.2 |
| 53 | VD24371 | 1 | 2021 | EFB symptomatic | negative | 34.9 |
| 54 | VD24740 | 1 | 2021 | EFB symptomatic | negative | 39.5 |
| 55 | VD25749-1 | 1 | 2021 | EFB symptomatic | negative | 19.5 |
| 56 | VD25749-2 | 1 | 2021 | EFB symptomatic | negative | 31.3 |
| 57 | VD25754 | 1 | 2021 | EFB symptomatic | negative | 18.7 |
| 58 | VD25791 | 1 | 2021 | EFB symptomatic | 34.6 | 18.5 |
| 59 | VD26510 | 1 | 2021 | EFB symptomatic | negative | 35 |
| 60 | VD27370 | 1 | 2021 | EFB symptomatic | negative | 21.4 |
| 61 | VD27554 | 1 | 2021 | EFB symptomatic | negative | 39.3 |
| 62 | VD28080 | 1 | 2021 | EFB symptomatic | negative | 22.7 |
| 63 | VD29139 | 1 | 2021 | EFB symptomatic | negative | 36.3 |
| 64 | VD31648 | 1 | 2021 | EFB symptomatic | negative | 22.1 |
| 65 | VD33548 | 1 | 2021 | EFB symptomatic | negative | 19.9 |
| 66 | VD33766 | 1 | 2021 | EFB symptomatic | negative | 38.4 |
| 67 | VD33909 | 1 | 2021 | EFB symptomatic | negative | 34.3 |
| 68 | VD33939 | 1 | 2021 | EFB symptomatic | negative | 38.7 |
| 69 | VD34400 | 1 | 2021 | EFB symptomatic | negative | 33.9 |
| 71 | VD34423 | 1 | 2021 | EFB symptomatic | negative | 36 |
| 72 | VD35085 | 1 | 2021 | EFB symptomatic | negative | 39.3 |
| 73 | VD35681 | 1 | 2021 | EFB symptomatic | negative | 36.9 |

**Table S8.** Pairwise comparisons by Log-Rank test comparing the survival between brood inoculated with 10^3^, 10^4^ and 10^5^ *Paenibacillus melissococcoides* 3.2 and a non-inoculated control. The Bonferroni-Holm P-value adjustment method for multiple comparisons was used.

|  | **control** | **10^5^** | **10^4^** |
| --- | --- | --- | --- |
| **10^5^** | 1.8e-12 |  |  |
| **10^4^** | 3.1e-05 | 0.0002 |  |
| **10^3^** | 0.0011 | 2.7e-06 | 0.21213 |

**Table S9.** Survival pairwise comparisons by Log-Rank test between one-, three- and five-day old inoculated brood with *Paenibacillus melissococcoides* 3.2 and non-inoculated control. The Bonferroni-Holm P-value adjustment method for multiple comparisons was used.

|  | **control** | **day 1** | **day 3** |
| --- | --- | --- | --- |
| **day 1** | 1.8e-12 |  |  |
| **day 3** | 4.1e-08 | 0.00657 |  |
| **day 5** | 1.6e-05 | 0.00031 | 0.18171 |

**Table S10.** Primers and probes used in qPCR method for the detection of *Paenibacillus melissococcoides* and *Melissococcus plutonius*.

| Organism | Gene | Primer/Probe | Oligonucleotide sequence (5’-3’) | References |
| --- | --- | --- | --- | --- |
| *P. melissococcoides* | sodA | PM-F | TCA TGG CGG AGG GCA TT | This study |
|  |  | PM-R | CCG TCG GGC TCA TGA TG |  |
|  |  | PM-Probe | FAM - CGG GCA TAG CCT G – MGB-Q500 |  |
| *M. plutonius* | napA | MP-F | GAC CTG TTT AGC TAT TAT CAC TA | Dainat et al., 2018 |
|  |  | MP-R | CAC CTA CAA TGA ATG ATT CAT TC |  |
|  |  | MP-Probe | FAM - TCC GCC TAA GCT ACC ACC TAA GAA C – BHQ1 |  |
